# Improved Injury Detection Through Harmonizing Multi-Site Neuroimaging Data after Experimental TBI: A Translational Outcomes Project in NeuroTrauma (TOP-NT) Consortium Study

**DOI:** 10.1101/2025.04.15.649026

**Authors:** G. Kislik, R. Fox, A.V. Korotcov, J. Zhou, M. Febo, Babak Moghadas, Adnan Bibic, Yunfan Zou, Jieru Wan, R.C. Koehler, T. Adebayo, M.P. Burns, J.T. McCabe, K.K. Wang, J.R. Huie, A.R. Ferguson, A. Paydar, I.B. Wanner, N.G. Harris, the TOP-NT Investigators

## Abstract

Multi-site neuroimaging studies have become increasingly common in order to generate larger samples of reproducible data to answer questions associated with smaller effect sizes. The data harmonization model NeuroCombat has been shown to remove site effects introduced by differences in site-related technical variance while maintaining group differences, yet its effect on improving statistical power in pre-clinical models of CNS disease is unclear. The present study examined fractional anisotropy data computed from diffusion weighted imaging data at 3 and 30 days post-controlled cortical impact injury from 184 adult rats across four sites as part of the Translational-Outcome-Project-in-Neurotrauma (TOP-NT) Consortium. Findings confirmed prior clinical reports that NeuroCombat fails to remove site effects in data containing a high proportion-of-outliers (>5%) and skewness, which introduced significant variation in non-outlier sites. After removal of one outlier site and harmonization using a global sham population, harmonization displayed an increase in effect size in data that displayed group level effects (p<0.01) in both univariate and voxel-level volumes of pathology. This was characterized by movement toward similar distributions in voxel measurements (Kolmogorov-Smirnov p<<0.001 to >0.01) and statistical power increases within the ipsilateral cortex. Harmonization improved statistical power and frequency of significant differences in areas with existing group differences, thus improving the ability to detect regions affected by injury rather than by other confounds. These findings indicate the utility of NeuroCombat in reproducible data collection, where biological differences can be accurately revealed to allow for greater reliability in multi-site neuroimaging studies.

**Significance Statement:** This project demonstrates the utility of NeuroCombat in reducing site effects in multi-site rodent imaging. We also demonstrate that harmonization improves the ability to distinguish between sham and injured rats at the voxel level and increase statistical power and effect size in areas of injury. Multi-center studies are becoming more common to allow for increased efficiency in data collection, and with conservative approaches and analysis into the datasets, NeuroCombat can be utilized to improve study reliability and reproducibility.

## Introduction

Diffusion-weighted imaging (DWI) is frequently used to study restricted water movement in brain tissue, and has been shown to be especially useful for investigating white matter pathology after traumatic brain injury (TBI) (1–3). Fitting DWI data to the tensor produces several metrics which describe the motion and direction of fluid, including the ratio of the primary eigenvalues- fractional anisotropy (FA) (4). Clinical multi-site studies using DWI are increasingly common, largely since the increased sample size results in improved statistical power (5,6). New multi-center, preclinical studies are also beginning to be published (7,8) so that it will be important to determine whether the application of tools and algorithms that are currently being used for clinical data harmonization can also be used preclinically. An important component of such multi-center studies is that they afford greater sensitivity to detect group differences, allowing hypotheses to be tested that have smaller effect sizes. These effects would otherwise not be detected because no single laboratory could easily produce the large sample size required. However, an unwanted consequence of combining data from multiple sites is the technical or non-biological variation that is introduced as a result of differences in equipment, especially scanner vendor and protocol type, among other factors (9). This undesired variation is referred to as site effects and has been a key target to improve the reproducibility of multi-site neuroimaging studies.

The development of multi-site studies has also come with an enhanced need to ensure data access and reproducibility. Noted in the FAIR principles which promote findability, accessibility, interoperability, and reusability, data integration both within and outside of consortia is key to ensuring good stewardship and accessibility to information for all stakeholders of a project (10). The growing size of data collected presents a further challenge for multi-site studies. Fluctuations in procedure, data analysis and collection, as well as instrumentation that result in site effects, can hinder replicability and the ability to compare data. When prospective alignment of procedures before data acquisition fails to enhance data harmonization, it becomes necessary to use retroactive methods to remove non-biological variation from data.

Numerous statistical methods have been created to harmonize neuroimaging data from multiple sites in order to remove site effects but preserve group differences (11). NeuroComBat was proposed (12) as a fork from ComBat which was initially developed to remove batch effects in genomic data (13). NeuroComBat (ComBat) has been shown to be able to consistently remove inter-site variation using an empirical Bayesian approach (12,13). It has been employed to successfully remove site effects across several research locations and clinical protocols without obscuring biological variation (14,15). It is notable, however, that in the absence of site effects, ComBat paradoxically *reduced* detectable biological variation among clinical data (16) indicating that some caution is required when applying this methodology.

While a large number of studies have demonstrated the effectiveness of ComBat in clinical populations, few have applied it to neuroimaging data from preclinical models of TBI. The Epilepsy Bioinformatics Study for Anti-epileptogenic Therapy (EpiBios4Rx) preclinical team previously demonstrated removal of magnetic field strength effects on FA values in the corpus callosum following ComBat harmonization, and this revealed significantly reduced FA in TBI rats (17). In that work, consistent areas of injury were demonstrated from data acquired across different scanners at the same site. Herein we were able to corroborate this both across the four different research sites that were included in the Translational Outcome Project in Neurotrauma (TOP-NT), as well as across scanners with different field strengths. Unlike in previous work where the injuries were consistent across scanners (17), the severity of injury used in the current study was purposely varied to test for the utility of FA and other tensor-derived dependent variables as potential biomarkers predictive of outcome. We therefore took injury severity into account when creating the ComBat model, using the degree of whole brain atrophy as a proxy. Additionally, previous studies have demonstrated increased statistical power as a result of harmonization, but have not examined voxel-level change in order to determine the potential effects on a viable injury site versus more remote regions. Thus, in addition to assessing whole-brain level univariate volumes of pathology across groups, we also assessed ComBat-related changes in statistical power, effect size and variability at the whole-brain level across individual *voxels* between groups. These measures, in conjunction with univariate harmonization, allowed us to investigate the ideal level at which ComBat harmonization could be applied to preclinical brain injury neuroimaging data, as well as to evaluate brain regional changes as a result of harmonization.

We found that NeuroCombat harmonization does decrease site-specific effect in univariate and voxel-wise measures of FA, thus better delineating biological differences as a result of injury. Analysis further displayed improved harmonization with the use of a pooled sham population and removal of significant outliers.

## Methods

### Experimental Protocol

As part of the TOP-NT consortium project, there were four data acquisition sites: The University of California Los Angeles (UCLA), the University of Florida and Morehouse School of Medicine (UF/MSM), Georgetown University and Uniformed Services University (GU/USU), and John Hopkins University (JHU), and a data analytics site - University of California San Francisco (UCSF). All study protocols used were in compliance with the Public Health Service Policy on Humane Care and Use of Laboratory Animals and approved by each site-specific Animal Research Committee, UCLA Approval #2021-082; UF/MSM Approval # 202010145; GU/USU Approval APG-21-044; JHU Approval #RA22M370. A total of 184 adult male and female, Sprague Dawley rats were used for this study and were randomly assigned to each group (UCLA: 33 TBI, 17 sham, JHU: 33 TBI, 12 sham, GU/USU: 33 TBI, 16 sham, UF/MSM: 26 TBI, 8 sham). Diffusion imaging data were acquired at 3 and 30 days after either controlled cortical impact (CCI) injury or sham anesthetic controls in adult, male (Mean<u>±</u>SD body weight 317±43.7g, 271.3±20.3g, 225±14.8g, 369±16.1g, at UCLA, GU/USU, JHU and UF/MSM, respectively) and female rats (body weight 254±19.1g, 239.3±13.8g, 183±12.7g, 256±7.9g, at UCLA, GU/USU, JHU and UF/MSM, respectively). A total of 343 scans were collected across the two post-injury times across all sites (25 datasets were missing). Prior to data acquisition, the injury and imaging protocols were harmonized across the sites using standard operating procedures (7,18). Post-acquisition data analysis, including all pre and post-processing data steps were also harmonized using a BASH script that wrapped multiple commands which was shared across the sites. The detailed methods below document the cross-site harmonized methods and any differences across sites are noted. All data were acquired with adherence to the ARRIVE guidelines (19) including group assignment randomization, blinding by use of unique animal IDs across site with semi-automated analysis routines, as well as conforming to the other eight guidelines on protocol, analysis and reporting standards.

### Controlled Cortical Impact

Rats were housed in pairs before and after the surgery in standard cages with ad libitum access to water and food. Rats were maintained on a 12:12 hour light/dark cycle in the vivarium. Estrus cycle was assessed for female rats by vaginal lavage/swab at the time of surgery to determine the stage of the estrus cycle of the rat on the injury day. Two different levels of CCI were induced at each site by varying the deformation depth below the dura of the impactor using similar methods across the sites (**Table 1**). Rats were anesthetized with 5% isoflurane vaporized in 1 L/min O2 (2% isoflurane was used for maintenance of the anesthesia with adjustments between 1.5 and 2%, and then were head-fixed in a stereotactic frame and their body temperature was maintained at 37°c using a temperature-controlled heating-pad. After prophylactic administration of analgesics according to the site-specific animal protocol, aseptic technique was used to make a midline, longitudinal skin incision, followed by a 6mm craniectomy (5mm UF/MSM) centered at +3.5mm anteroposterior and 3mm lateral to the midline over the left hemisphere using a dental drill cooled under intermittent saline. CCI injury was conducted using a 5mm (4mm UF/MSM) diameter, bevel-edge, metal impactor tip at an angle parallel to the dural surface and at impact depth below dura, speeds and dwell times ranging from: 1.3-2.3mm, 3.5-6m/s and 100-240ms, using an electromagnetic impact system (Leica Biosystems, IL, USA). The craniectomy was closed with a non-toxic sealant (Kwik-Cast, WP instruments, USA) and the skin sutured.

**Table 1:** Injury Parameters Across the Sites.

|  | **UCLA** | **GT/USU** | **JHU** | **UF/MSM** |
| --- | --- | --- | --- | --- |
| **Craniectomy Size (mm)** | 6 | 6 to 7 | 6 | 4 |
| **Antero-Posterior Position (mm)** | -3.5 | -3.5 | -3.5 | -2.5 |
| **Medial-Lateral Position (mm)** | 3.5 | 3.0 | 3.0 | 3.0 |
| **Hemisphere** | left | left | left | right |
| **Tip Diameter (mm)** | 5 | 5 | 5 | 4 |
| **Injury Device** | Leica One | Leica One | Leica One | Leica One |
| **Tip Material** | metal | metal | metal | metal |
| **Tip Shape** | flat | round | flat | flat |
| **Angle of Impactor (degrees)** | 13-18 | 12 | 5-10 | 0 |
| **Dwell time (ms)** | 200 | 200 | 200 | 200 |
| **Speed (m/s)** | 5 | 5 | 5 | 5 |
| **Depth Below Dura (mm)** | 1.7; 2.1 | 1.7; 2.1; 2.8 | 1.0; 1.2; 1.5; 1.7 | 1.7; 2.1 |
| **Angle to Perpendicular (degrees)** | 10-15 | 5-10 | 5-10 | 0 |
| **Avg. Time Under Anaesthesia (means ± STD; min)** | 39 ±0.01 | 45.8±12.7 | 25.1±1.5 | - |

### Image Acquisition

Data were acquired on Bruker Biospec consoles (Bruker, Billerica, MA, USA) connected to magnets from a variety of vendors and with different field strengths (Tesla, T) across the sites (7T -UCLA and GU/USU, 11.7T – JHU and 11.1T - UF/MSM) with gradients of a varying peak strength and rise time resulting in different slew rates. Radiofrequency coils were similar across three sites (birdcage volume transmit coil decoupled from a receive-only 4-channel phased-array surface coil) and a transceive quadrature coil at UF/MSM. Rats were imaged prone under isoflurane sedation (1-1.5%) vaporized in medical air (0.5l/min) positioned in a cradle with head stabilization using a 3-point head fixation. The diffusion data were acquired after a number of other scans within the same imaging session as part of a larger protocol, taking around 60mins prior to diffusion imaging. The prior scans for functional and structural imaging were conducted under a mixture of continuous dexmedetomidine sedation (0.05mg/kg/hr, subcutaneously) and isoflurane (0.5%) prior to switching to isoflurane alone. Temperature was maintained at 37°C via homeothermic-controlled external air or water heating.

A 3-dimensional, single-shot, spin echo, echo planar imaging sequence was used to acquire diffusion-weighted images with directionally-encoded, monopolar diffusion gradients applied along 88 different, non-colinear directions, with b values of 0, 1000 and 3000 s/mm^2^ (n=4, 44 and 44, respectively), and no averaging. Four left-right, phase-reversed, b=0 images were also acquired. Repetition and echo times varied slightly according to the supported gradient hardware (**Table 2)**. Similarly, the diffusion gradient amplitude (big delta) was varied across site to arrive at similar b values while maintaining similar diffusion times, the time between application of the diffusion-sensitizing gradients (**Table 2**). The acquisition data matrix was constrained to a 72x49x96 matrix size, in the 1^st^ phase-encoding, 2^nd^ phase-encoding and read-out directions, within a field of view (FOV) of 18x12.5x24mm, respectively in the dorsal-ventral, left-right and antero-posterior planes, resulting in an isotropic resolution of 250µm. Outer-volume signal suppression using multiple, 5-10mm saturation slices were applied to reduce signal from outside the brain.

**Table 2:** MRI Parameters for DWI and MGE Parameter sequences Across the Sites. <u>Key</u> SE-EPI= spin echo, echo-planar imaging.

| **DWI** | UCLA (7T) | GU/USU (7T) | JHU (11.7T) | UF/MSM (11.1T) |
| --- | --- | --- | --- | --- |
| **Mode** | 3D-SE-EPI, SingleShot | 3D-SE-EPI, SingleShot | 3D-SE-EPI, SingleShot | 3D-SE-EPI, SingleShot |
| **FOV (mm)** | 24x18x12.5 | 24x18x12.25 | 24x18x12.25 | 24x18x12.25 |
| **Matrix Size (#voxels)** | 96x72x49 | 96x72x49 | 96x72x49 | 96x72x49 |
| **Resolution (um)** | 250 | 250 | 250 | 250 |
| **TR (ms)** | 1000 | 1000 | 1000 | 1000 |
| **TE (ms)** | 28.4 | 30 | 27.85 | 27.85 |
| **Flip Angle (degrees)** | 90 | 90 | 90 | 90 |
| **BandWidth (KHz)** | 357 | 300 | 357 | 357 |
| **Big Delta (ms)** | 12 | 12 | 12 | 12 |
| **LittleDelta (ms)** | 5.3 | 5 | 5.35 | 5.35 |
| **Bvalue1 (S/mm<sup>2</sup>)** | 1000 | 1000 | 1000 | 1000 |
| **Bvalue2 (S/mm<sup>2</sup>)** | 2800 | 3000 | 3000 | 3000 |
| **MGE** |  |  |  |  |
| **Mode** | 3D | 3D | 3D | 3D |
| **FOV (mm)** | 25.6x19.52x12.8 | 32x32x16 | 25.6x25.6x19.2 | 25.6x25.6x19.2 |
| **Matrix Size (#voxels)** | 160x122x80 | 200x200x100 | 160x160x120 | 160x160x120 |
| **Resolution (um)** | 160 | 160 | 160 | 160 |
| **TR (ms)** | 125 | 45 | 45 | 45 |
| **TE Range (ms)** | 2.8-51.8 | 2.5-26.5 | 2.5-26.5 | 2.5-26.5 |
| **Echo Spacing (ms)** | 4.9 | 3 | 3 | 3 |
| **Number Echoes** | 11 | 9 | 9 | 9 |
| **Flip Angle (degrees)** | 20 | 13 | 13 | 13 |
| **BandWidth (KHz)** | 100 | 100 | 100 | 100 |

Whole brain anatomical data were acquired using a 3D-multi-gradient echo sequence with a variable matrix size and field-of-view across site, resulting in the same isotropic resolution of 160µm (**Table 2**). Eleven echoes were acquired from 2.8-51.8ms with an echo spacing of 4.9ms, a repetition time of 45-125ms and a 13- 20 flip angle (**Table 2**).

### Image Preprocessing

Bruker raw data were converted to nifti (20), brain extracted (21), denoised (22), and corrected for Gibbs ringing (23), after which phase distortions were unwrapped with TOPUP (24,25) and Eddycorrect (26), all of which were implemented under MRTRIX (27). Data were then fit to the tensor to derive scalar images- fractional anisotropy (FA), axial, radial and mean diffusivity (AD, RD, MD, respectively). FA data were then used to derive an unbiased mean deformation template (MDT) at each site using successive registrations of rigid, affine and non-linear alignment under ANTS (28). The resulting affine transformations and warp fields were applied to all scalar data to align them to the local site MDT. Voxel-wise sham mean and standard deviation (SD) maps were computed for each tensor indices, which were then used together with each co-registered brain volume scalar to calculate regions of high and low indices relative to local sham data at a corrected z-score threshold based on a distribution-corrected z-score (29, 42) beginning with a desired z value of |z|>±3.1 (p<0.001). The resulting corrected z values for thresholding the injury groups at each site at 03d/30d was: 3.85/3.75, 3.99/3.85, 4.19/4.19 and 5.08/5.08, respectively at UCLA, GU/USU, JHU, UF/MSM. For the sham group the adjusted z values at 03d/30d was: 2.67/2.72, 2.61/2.67, 2.52/2.52 and 2.38/2.38, respectively at UCLA, GU/USU, JHU, UF/MSM. The brain-wide volumes of tissue surviving these upper and lower z-score thresholds for each tensor scalar, herein designated Scalar_LOW_ and Scalar_HIGH_ volumes, were then quantified for each brain by counting the voxels that were greater or less than the z-score. For voxel-level harmonization, a multi-site MDT was calculated using the site-specific MDTs as input, and then each site-specific subject space data were transformed to the mean site MDT in a single step through application of the original warp and affine transformation together with those generated from site MDT to multi-site MDT space. Calculation of univariate data within this common space occurred using the same corrected z-scores used for univariate level harmonization within site-specific space.

### Tensor-Based Deformation Analysis

Anatomical data were used to assess local tissue deformations to approximate tissue atrophy/compression and swelling/expansion as we have before (30,31). Briefly, a mean deformation brain template was constructed from all data using ANTS (32) and the Jacobian Determinant was calculated from the resulting warp transformation fields. Sham Jacobian data were used to calculate voxel-based mean and standard deviation maps which were then used to calculate z statistic maps for all injured data. The volume of tissue surviving a threshold of z ± 3.01 (P < 0.01, uncorrected) was determined for each injured brain and used to assess the region of local tissue contraction, and referred to herein as tissue atrophy.

### Univariate and Voxel-Level Harmonization

The R package neuroCombat_1.0.13 was used for harmonization (https://github.com/Jfortin1/neuroCombat_Rpackage). The covariates included in the model were day post-injury, site, atrophy, and sex. All scans across days and groups were combined within the NeuroCombat model to generate harmonized measurements. All statistical analysis was conducted in RStudio version 2023.6.0.421 (Posit Team 2023). Loading of NIFTI files and generation of whole-brain maps of changes in effect size, standard deviation and power were conducted with the oro.nifti and neurobase packages (33,34). Cohen’s d was calculated as a measure of effect size by dividing the difference between the mean injured and mean sham animal by the pooled standard deviation. Plotting of violin plots for univariate data was done using the ggplot2 package (35). The False Discovery Rate (FDR) correction was applied to correct for multiple comparisons. Data points for voxels exceeding the range of 1.5 times the interquartile range (IQR) from the first (Q1) or third (Q3) quartiles were identified as outliers. Four-site data was visualized using a residual plot with values predicted by a regression model generated from the harmonized data across each site, in order to demonstrate the presence of unequal variance among sites post-harmonization. FSLeyes was used to visualize effect size and power maps (36) and the Estimation Stats software package was used to visualize univariate effect sizes (37). Bland Altman plots were produced by plotting the difference of each paired voxel measurement against the two voxels’ average value for each combination of site. Frequency maps were generated by finding the number of rats different from mean sham at each voxel using a modified z-test (the quantity of the test value minus the sham mean at that voxel divided by the standard deviation of shams). Power at each voxel was calculated with the R pwr library’s pwr.t2n.test function (38). Relevant code to reproduce the analyses found in this study can be found at the following link: (https://github.com/ngharris/neurocombat_TBI)

## Results

For clarity, we limited the reporting of data harmonization to fractional anisotropy, but we have included data on the other tensor scalers MD, AD and RD within the supplementary section (**Supplementary Figs. S1-15**). These data were in general, qualitatively similar to the FA results described herein. Volume of atrophy was included as a covariate in harmonization to account for intentional variation in injury severity, but these volumes were not significantly different between sites (one-way ANOVA, p>0.05).

The number of brain voxels with FA greater or less than a statistical theory-derived distribution-corrected z-score (equivalent to P<0.001 relative to site-specific sham data, Fig. 1) are referred to as FA_High_ and FA_Low_ volumes of pathology, respectively. We have operationalized “pathology” based upon a voxel tensor scalar value being lower or greater than the z-score threshold. It should be noted that “pathology” in shams is not due to injury, but other likely sources of variance, such as between scanner effects and animal-to-animal differences. Two harmonization methods were used. In the first, the volume of whole-brain pathology was derived from harmonization of univariate volume data derived from subject data co-registered to site-specific brain template space, and is herein referred to as univariate-level harmonization. In the second method, the volume of pathology was derived after harmonization of whole-brain FA data at the brain voxel level, derived from data coregistered to a multi-site-derived template, and is herein referred to as voxel-level harmonization.

**Fig. 1.**
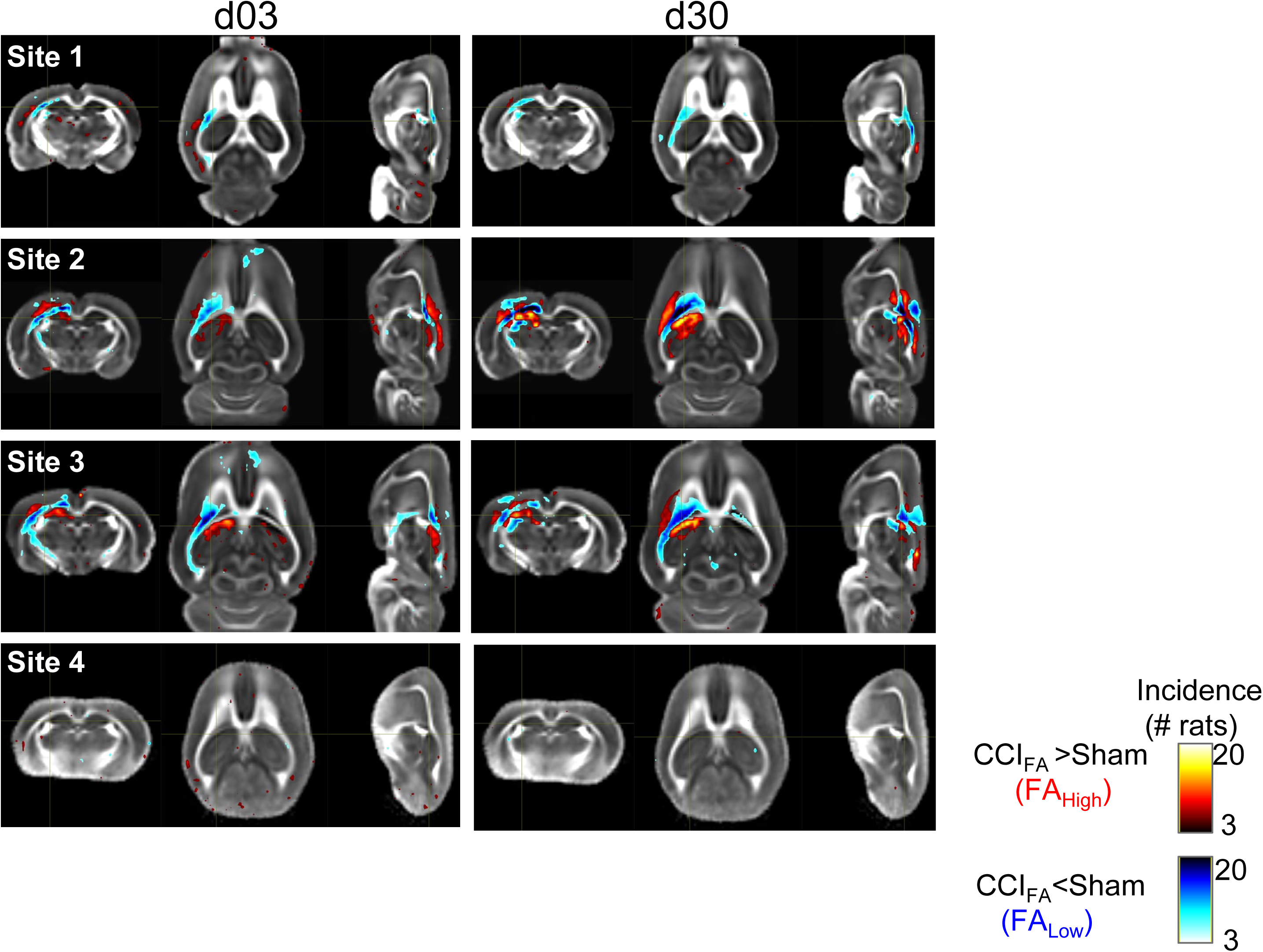
Voxel overlap maps of regions of FA-related pathology prior to harmonization at each site at three and 30d post-injury. Individual FA volume data from each injured rat that survived below the <u>FA_Low_</u> or above the FA<u>_High_</u> corrected zmap-threshold (red, blue colors respectively, P<0.001) relative to local sham data are plotted as binary incidence maps for each site at day 3 and 30 post-injury. The voxel pseudocolors represent the overlap between the number of rats where FA survived threshold at each brain voxel location. Site 4 shows low overlap between rats.

### Outlier Sites Lead to Increased Variance in Non-Outlier Sites After Univariate-Level Harmonization

Previous work (39) has shown that if the proportion of outliers is higher than 5% of the data, or if extreme outliers exist, this can severely distort the ability of NeuroCombat to remove site effects. In the current data, we found that the variance for both high and low FA values was significantly increased within Site 4 sham data as compared to all other sites for univariate data before harmonization (P<0.001). This increased variance within site 4 data was maintained even after harmonization when compared to the other three sites (p<0.001). Confirming these data, the residuals of FA values for sham rats predicted by a regression model generated with harmonized sham data showed unequal distributions with the largest spread in site 4 data. One potential cause of this heteroscedasticity could be attributed to the presence of outliers, which could lead to ineffective harmonization if all sites were included. (**Fig. 2**). However, log transformation of these data revealed similar results with a greater number of univariate outliers in site 4 compared to all and significantly increased variance. Removal of site 4 data resulted in an absence of site effects in the harmonized data (P>0.05), further substantiating these findings. As a result, all further univariate analyses were conducted with data from only three sites in order to more accurately determine the utility of using NeuroCombat to harmonize preclinical imaging data from multiple sites. Additional analyses using data from all four sites can be found in supplementary data (**Figs. S16-18**).

**Fig 2.**
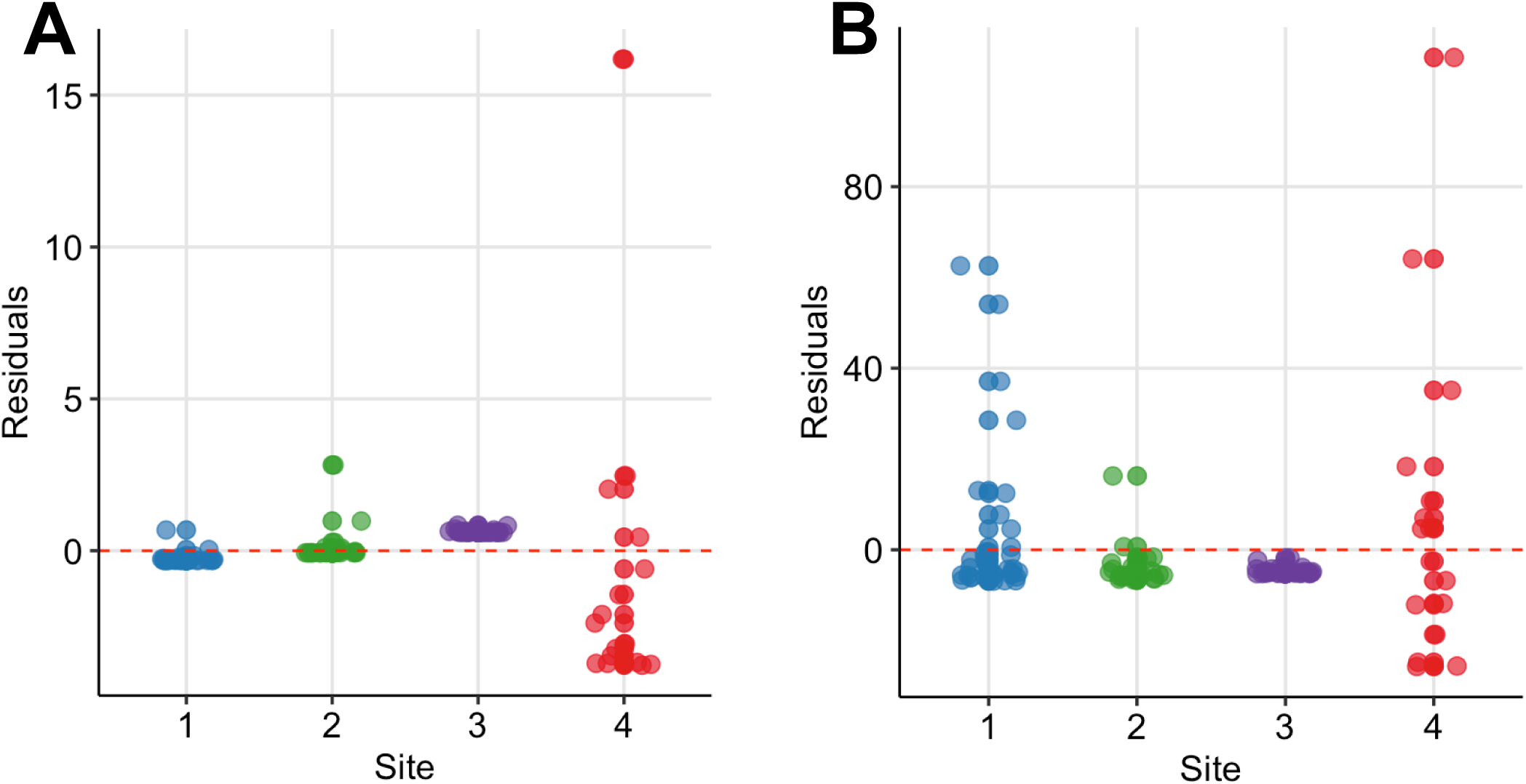
Impact of using all four sites for univariate-level harmonization. The residuals of (**A**) FA_Low_ and (**B**) FA_High_ predicted using a model generated from univariate 4-site NeuroCombat data demonstrates a larger spread of residuals in site 4 and with fewer points centered around 0.

### Univariate-Level Harmonization with Pooled but Not Site-Specific Shams Leads to Improved Detection of Injury

An important consideration when investigating differences in multi-site data is whether to pool shams from all sites, or whether to treat site-specific sham data independently for the calculation of univariate volume data. Pooled shams were used to generate the z-scored FA_Low_ and FA_High_ values for each rodent prior to harmonization. To gauge how effectively differences between sham and TBI rats were detected when considering pooled sham data versus sham data specific to each site, we examined FA_Low_ and FA_High_ distributions across the sites. Compared to unharmonized data, harmonization with pooled shams led to a 32% increase in the number of injured rats across all sites, where FA_Low_ volumes of pathology were more than two standard deviations larger than the mean sham animal across all sites (n=21 vs 30 animals at day 3 and n=19 vs 23 at day 30, respectively). The number of injured rats with FA_High_ volumes above the mean after harmonization were reduced or remained similar to unharmonized data (n=9 vs 8 at day 3, respectively and no change at day 30).

Harmonization with site-specific sham data afforded an unexpected lower improvement compared to the prior reported pooled sham harmonization; the number of injured animals with FA_Low_ volumes above sham mean data after harmonization increased by only 10% (n=33 vs 32 at day 3 and n=52 vs 45 at day 30, respectively). However, the number of injured rats with significantly different FA_High_ volumes after harmonization increased by 17.6% (n=11 vs 9 at day 3 and n=9 vs 8 at day 30, respectively. While univariate-level harmonization led to mixed results overall, the greatest improvement occurred in injured rat FA_Low_ volumes using pooled sham data. As a result, since NeuroCombat is intended for use on data pooled from multiple sites, pooled shams were used for all further univariate analyses. Global shams were also used for voxel-level analyses for consistency.

### Group Differences after Univariate Data Harmonization Using Pooled Shams

Previous reports have demonstrated that removal of site-specific effects after harmonization would indicate that any remaining group differences could accurately be attributed to a true difference in pathology, thus demonstrating successful harmonization (12). Within the original, unharmonized, univariate data, there were significant sham vs TBI group effects at all time points for FA_Low_ volume (**Fig. 3A,C**, p<0.01 and p<0.001, respectively for 3 and 30d post-injury, linear mixed model ANOVA) and significant site effects for FA_High_ volume at both time points (p<0.001, **Fig. 3B,D**). However, after univariate-level harmonization, the group difference in FA_Low_ volume originally found at day 3 was lost (p>0.05). Furthermore, group differences were introduced at FA_High_ volumes at day 3 through harmonization (p<0.05). Finally, univariate-level harmonization failed to correct for the site effects observed in FA_High_ volumes at day 3 and 30 post-injury before harmonization (**Fig. 3B,D**) where sham data from Site 1 had substantially higher volumes of pathology and were more variable than the other two sites (p < 0.001, linear mixed model ANOVA). There was a significant effect of sex in low FA at days 3 and 30 both before and after harmonization (p<0.05, linear mixed model ANOVA), indicating that group effects seen after harmonization were not erroneously introduced due to sex used as a covariate. This result was similarly found in the other measures, where the few site effects found were not removed, but group differences were maintained post-harmonization **(Figs. S1,6,11).**

**Fig. 3.**
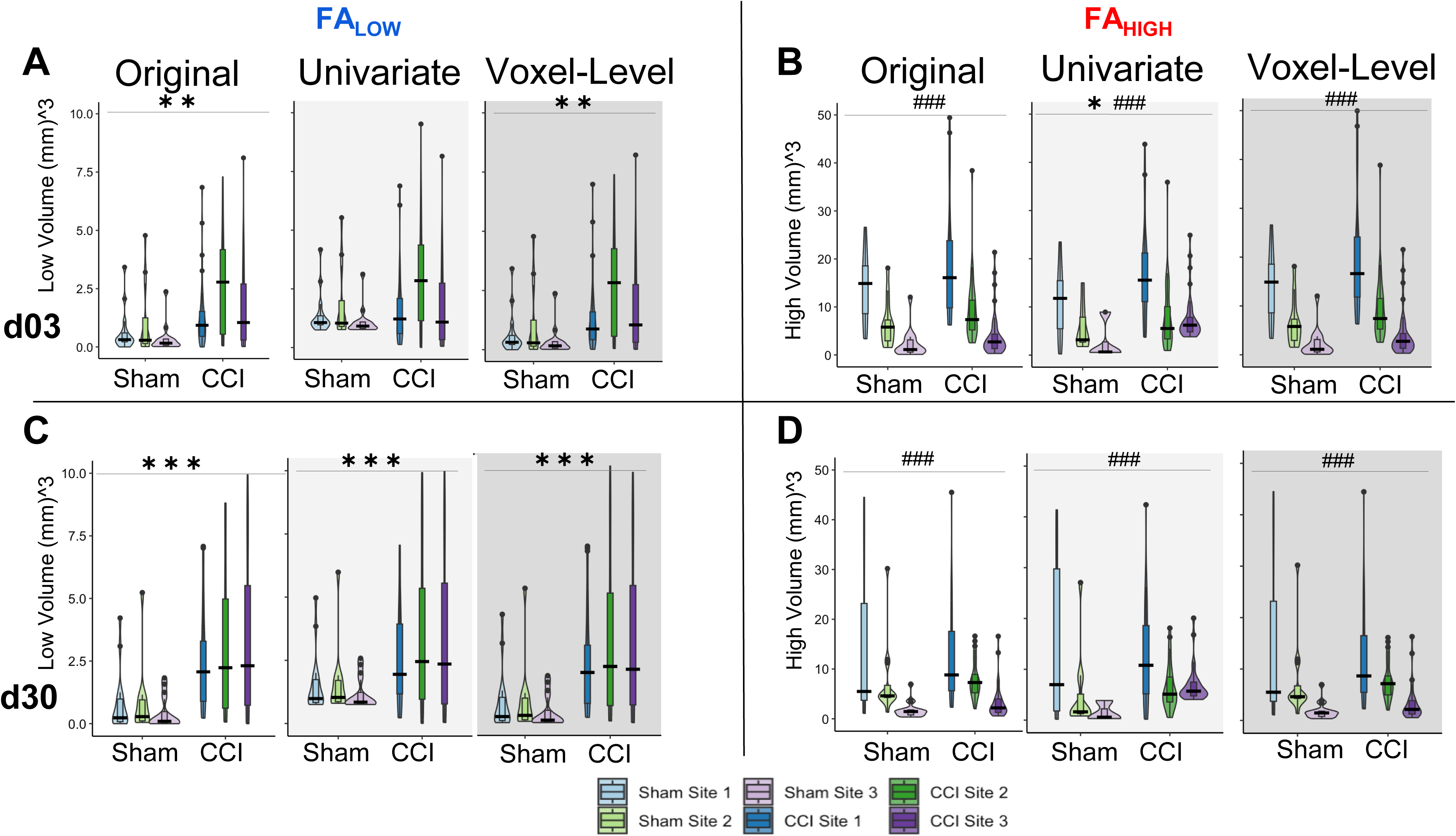
Effect of univariate and voxel-level harmonization on the burden of whole brain injury volume using pooled sham data. Brain volumes of pathology identified by (**A,C**) FA_Low_ and (**B,D**) FA_High_ at 3d (**A,B**) and 30d (**C,D**) are plotted for each site before (“Original”) and after univariate and voxel-level harmonization. ###/*** = a significant effect of location and injury, respectively, p<0.001, ##/** = p<0.01, #/* = p<0.05 (linear mixed model ANOVA).

Despite these unexpected effects, we calculated the effect size for each site in order to quantify any improvement in the ability to discriminate between experimental groups due to harmonization. Using the difference in univariate FA volume for injured versus pooled sham data at each timepoint, effect size changed in different directions for day 3 FA_Low_ data after harmonization at sites 1, 2 and 3 by -4.3, 1.35 and 2.1-fold respectively. This was significantly different from zero (p<0.05) for Sites 2 and 3 compared to pre-harmonization, when it was not previously different (**Fig. 4A**). Similar but smaller effects were found for day 30 data at all sites (**Fig. 4B**). Despite the introduction of group differences in the FA_High_ data at day 3 through harmonization, the magnitude of the effect size decreased for site 3 (p<0.05 pre-harmonization, p>0.05 after harmonization, **Fig. 4C**) and it remained similar for other sites (p>0.05).

**Fig 4.**
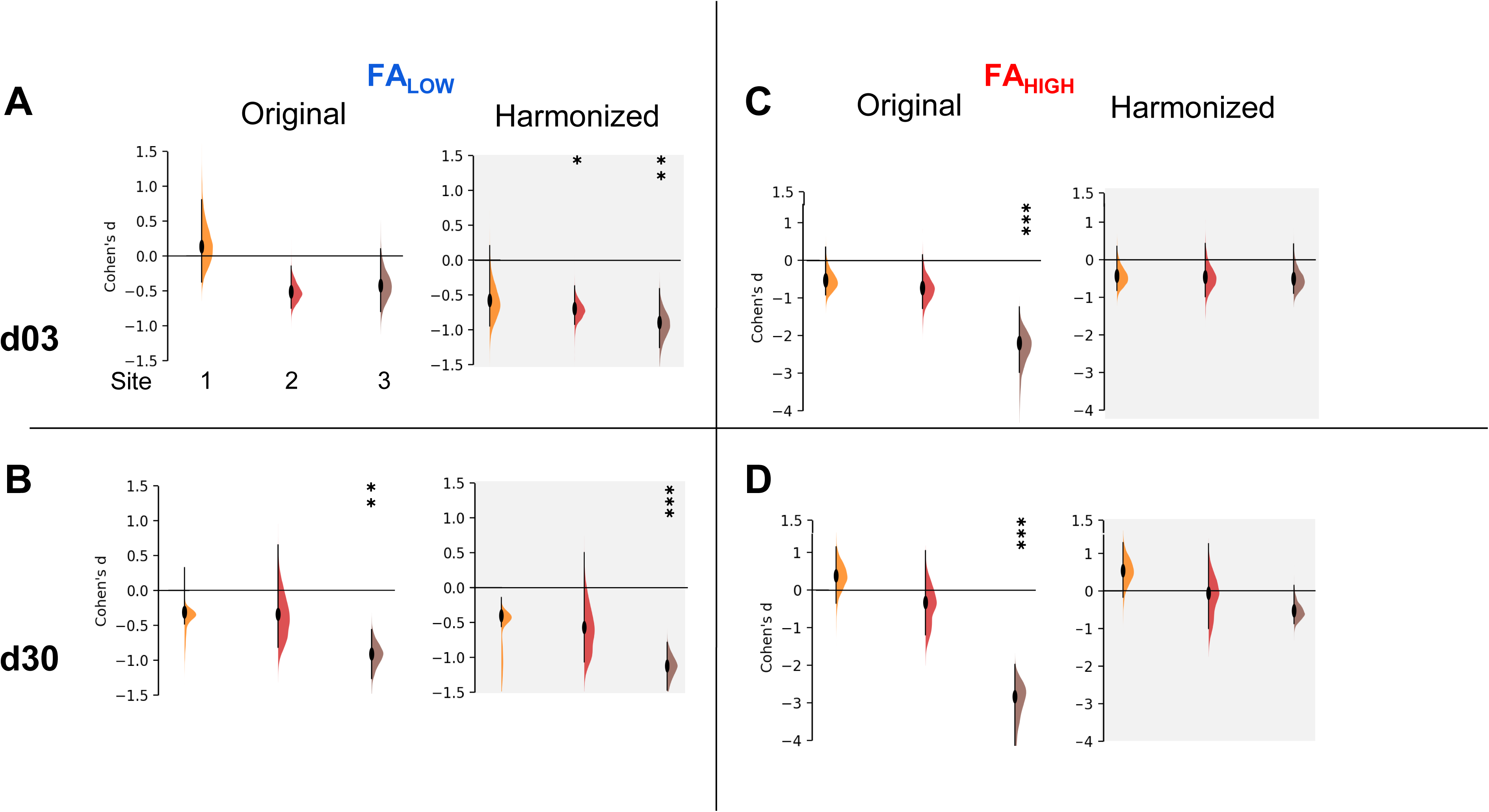
Effect sizes after univariate-level harmonization. Injury effect size measured by Cohen’s D for each site before and after univariate harmonization for (**A, B**) FA_Low_ volumes and (**C,D**) FA_High_ volumes at (**A,C**) d03 and (**B,D**) d30 post-injury. The data demonstrates an increased effect size for all sites for FA_Low_ volumes, at 3 and 30d (**A**, **B**) but no change or a removal of significant effect size across sites for FA_High_ volumes (**C**, **D**). *** = an effect size significantly greater than 0, p<0.001, ** = p<0.01, * = p<0.05 (t-test)

### Voxel-Level Harmonization Leads to a Common Distribution of FA Values and Effect Sizes Across Sites

By visual inspection of the group-level incidence maps of volumes of pathology described by FA_Low_ and FA_High_, we found there were marked differences at site 4 compared to Sites 1-3 at both 3 and 30 days after injury (**Fig. 1**), in agreement with the prior outlier volume analysis above. We confirmed this quantitatively for FA values; of the four sites analyzed, 10.5% of the FA values from Site 4 were found to lie more than 1.5 interquartile ranges outside the interquartile range of FA values when compared to the other three sites at each voxel location. The outliers among Sites 1, 2 and 3 were smaller, containing only 4.6%, 1.7% and 0.9% outliers, respectively, which was more than 2-fold less than Site 4. Additionally, Site 4 FA population data were found to have an excess kurtosis of 0.497, compared to -0.774, 0.159 and 0.274 in Sites 1, 2, and 3, respectively indicating a quite different distribution of FA volumes. These discrepancies were similar to the findings at the univariate volume level described in the prior sections. Harmonization of whole brain data resulted in large decreases in the standard deviation of Site 4 FA data and corresponding increases in variance in Sites 2 and 3, as well as to a lesser extent in Site 1 (**Fig 5A**). Since the presence of an outlier site severely biased the performance of NeuroCombat, further analysis of whole brain data was done with only Sites 1, 2 and 3.

**Fig 5.**
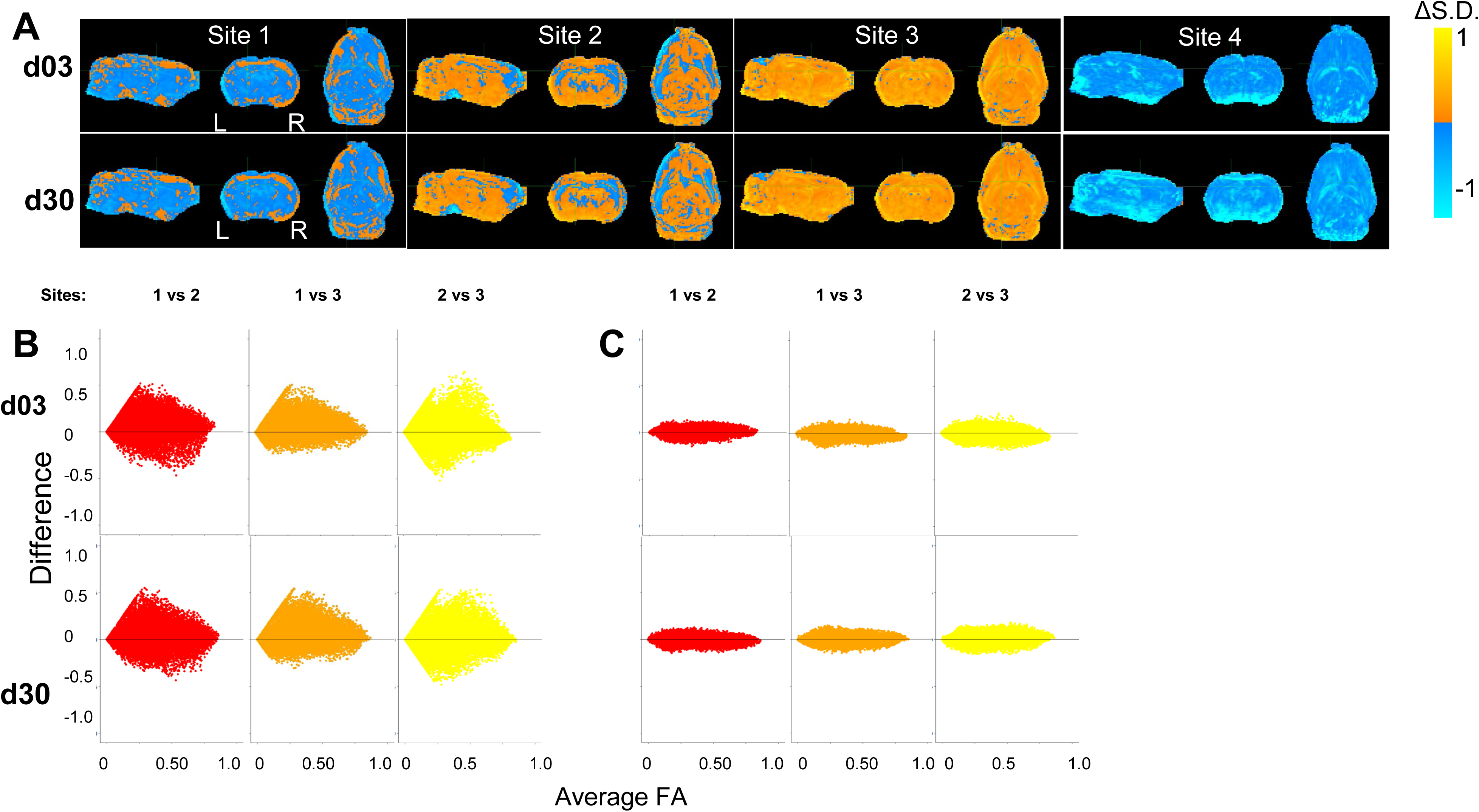
Harmonization at voxel-level resolution. **(A)** Change in standard deviation due to harmonization (harmonized – non-harmonized), where orange and blue voxels indicate an increase and decrease in standard deviations due to harmonization respectively. Site 4 shows a decrease in standard deviation across the brain, indicating a large variation in the original, non-harmonized sham distribution of FA values across the brain. Key: [L]=Left side of the brain, [R]=Right side of the brain. (**B-C**) Harmonization of FA values of each individual voxel across site pairs is demonstrated using Bland-Altman plots for (**A**) original, unharmonized values and (**B**) harmonized values. The difference between sites decreases due to harmonization.

Bland-Altman plots were used to quantify the effect of harmonization across the whole brain between sites at the voxel level across all subjects. These graphs plot the mean value of the same voxel location from each pair of sites against their difference, and they were generated for each pair of sites before and after harmonization. In the original, unharmonized data, large differences were observed for all combinations of sites indicating site-related differences. After NeuroCombat harmonization, the difference between voxel measures between pairs of sites approached zero, indicating that FA values were more similar at the same brain location across sites after harmonization. This was supported by a Kolmogorov-Smirnov analysis, which showed significantly different site effects before harmonization (p<0.001) and a resolution of these site effects after harmonization on average between each combination of sites (p>0.05, **Fig. 5B**).

Crucially, all group level effects shown prior to harmonization through univariate volume analysis were maintained using voxel-level harmonization: group difference in FA_Low_ at 3- and 30-days post injury were maintained (p<0.01), as well as site-specific differences in FA_High_ at 3- and 30-days post injury (p<0.01) (**Fig. 3**).

### Voxel-Level Harmonization Improves Group-Level Delineation of Injured Brain Regions and Leads to Spatially-Specific Statistical Improvements

Since the application of NeuroCombat at the voxel level resulted in decreased cross-site differences, confirming prior work at the same site across different field strengths (17), we next investigated the statistical effect of harmonization within regions of primary injury as well as in less injured regions across the brain. Correct interpretation of statistical results requires consideration not only of statistical significance but effect size and statistical power. Improvements in power, the probability of detecting a result when it is present, can help to detect significant differences in datasets with small effect sizes. Given the simple statistical design of the current study, increases in sample size from combining data in this study should generally be expected to increase statistical power at a geometric rate through reduction in standard error. However, power is also affected by effect size and unexplained variances, among other factors. Therefore, given the heterogeneity of the injury, due to technical variation of the primary insult and site-related effects of correctly observing the injury, both ideal sources of unexplained variance, we sought to determine how cross-site harmonization would affect both statistical power as well as the related effect size within different regions of the brain. We hypothesized that harmonization would increase the probability of correctly detecting group-level differences, particularly within specific brain regions that may exhibit high variability at the single site level through increases in power and effect size.

To accomplish this, whole brain images of change in power and effect size due to harmonization were generated by voxel-based subtraction between the original, non-harmonized and the harmonized FA data for each site (**Fig. 6**). The average effect from harmonization calculated over all three sites was an increase in power around the primary injury area of the ipsilateral corpus callosum, but decreases in most other regions (**Fig. 6A**). At the site level at day 3, power increased within the ipsilateral white matter and gray matter surrounding the primary injury area in all three site data, with the greatest improvement measured in Site 1 (**Fig. 6B**). There were, however, decreases in power as a result of harmonization, globally within the grey matter across all sites and at both times post-injury. At day 30, the greatest increase in power was observed in data from Site 1, similar to day 3, while there were power decreases in the striatum and ventral areas of the brain in data from Sites 2 and 3. These same remote changes also generalized to the AD, MD and RD (**Figs. S4,S9,S14**).

**Fig 6.**
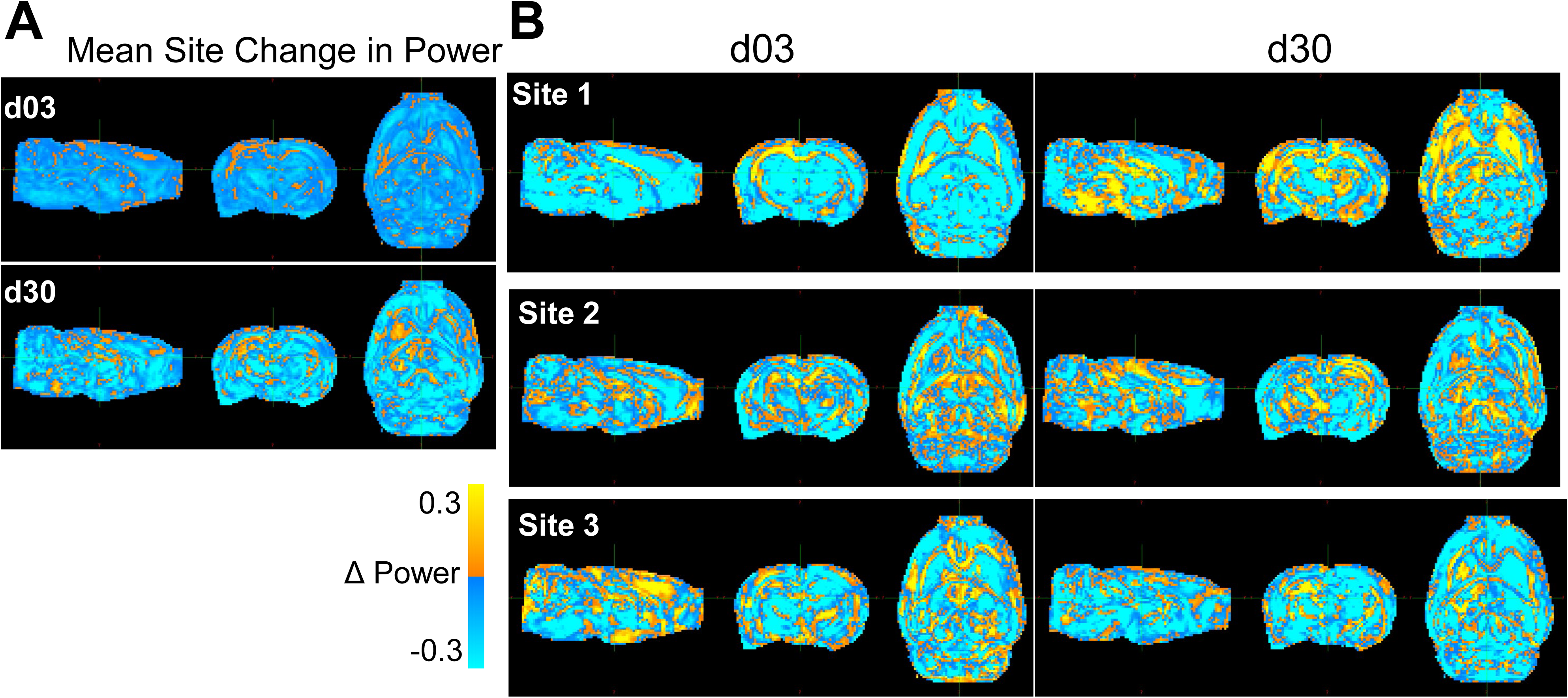
Whole-brain maps of regional changes in power due to harmonization. **(A)** Difference in power across a pooled average of all three sites post-injury due to harmonization and **(B)** across site and post-injury day. The magnitude scale -30% and 30% indicates the percent difference in statistical power between pre and post voxel- level harmonization, where yellow/blue indicate power increases and decreases, respectively due to harmonization.

Calculation of the corresponding voxel-wise change in effect size revealed an ipsilateral increase in the magnitude in the site-averaged data (**Fig. 7A**) consistent with the prior power changes (**Fig. 6A**). However, while power changes were decreased by harmonization throughout much of the brain, effect size increased over the majority of the brain remote from the primary injury site, with the exception of decreases within multiple clustered voxels that occurred at both post-injury times. Across site-averaged data, these trends persisted in AD, MD and RD (**Figs. S4,S9,S14**), but the drivers varied. For FA, the underlying data driving these site-wide effects were a decrease in effect size at site 1, but an increase at Sites 2 and 3 (**Fig. 7B**). The opposite trend was seen in MD and RD, where Site 1 showed increases in effect size in the injury area, while Sites 2 and 3 exhibited large decreases around the injury area (**Figs. S9 & S14**). AD showed a different trend, with Site 2 driving much of the decreases and Sites 1 and 3 showing increased effect size at the injury area (**Fig. S4**). The heterogeneity in both effect size and power across the sites outside of the primary injury area may be explained by the effect of NeuroCombat as it attempts to adjust for the variability introduced by site effects, ultimately converging at a common spatial pattern.

**Fig. 7.**
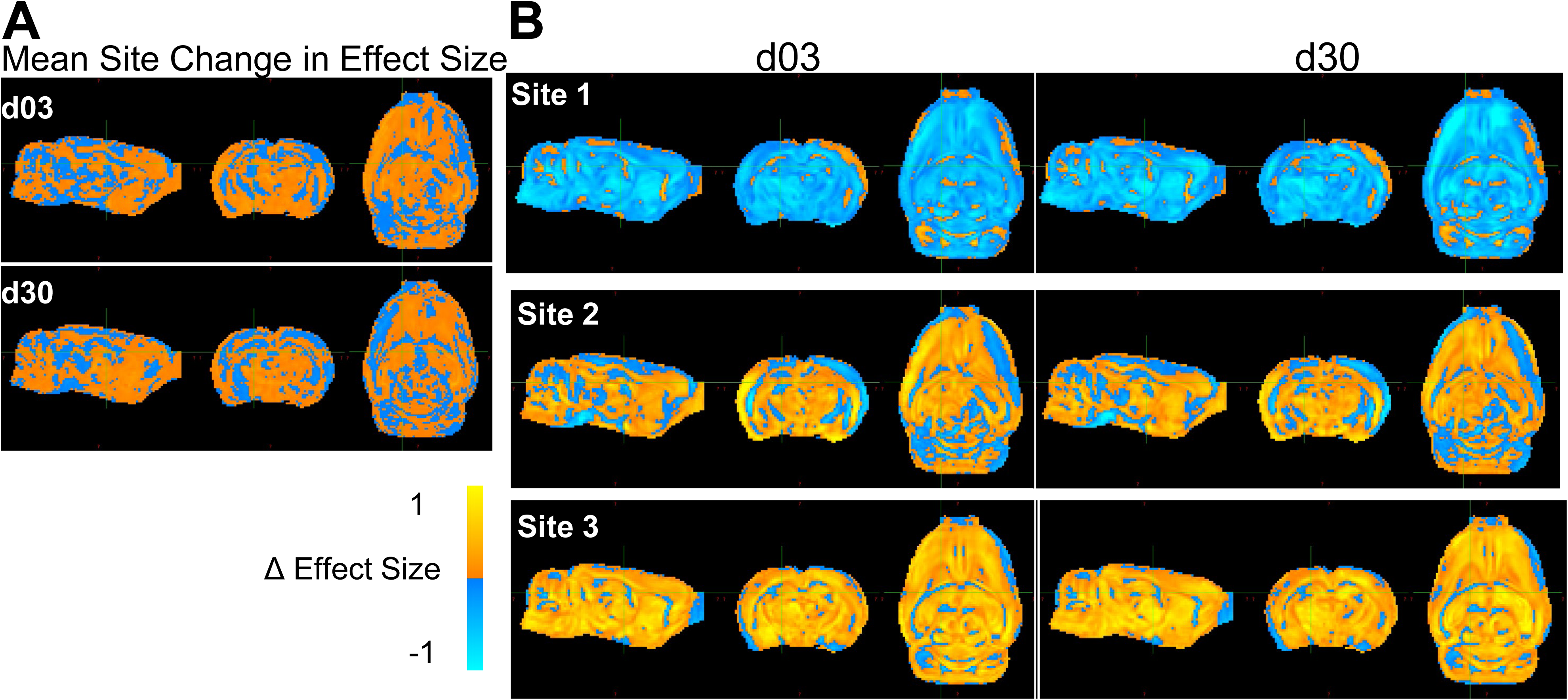
Whole-brain maps of regional changes in effect size due to harmonization. **(A)** Difference in effect size across a pooled average of all three sites post-injury due to harmonization and **(B)** across sites and post- injury day. The magnitude scale –1 and +1 Cohen’s d indicates the difference in effect size/voxel between pre and post voxel-level harmonization, where yellow/blue indicate increases and decreases, respectively due to harmonization.

We sought to determine whether the improved power and effect size was underpinned by the detection of pathologically low or high FA when compared to the sham group. FA voxel overlap maps showing the proportion of rats with FA values significantly different from sham after harmonization versus before, revealed an increased number, and overlap of rats after harmonization when compared to the original data (**Fig. 8**). These increases were especially prevalent within, and adjacent to, the primary, ipsilateral injury area. All differences detected were related to decreases in FA; no data was underpinned by increased areas of FA_High_ when compared to shams. These results indicate that cross-site data harmonization can improve the reliability of group differences in the areas surrounding the primary injury area at the cost of lower power in regions more remote from the primary injury site. For other scalar measures, results were markedly different. Generally, the proportion of rats significantly different from mean sham was lower. In the AD dataset, most areas of difference were found in the ipsilateral cortex, frontal pole and cerebellum. The only location to produce an increase in the number of rats was a small portion of the corpus callosum. In other areas, harmonization decreased the number of voxels with significant differences, suggesting an initially weak effect at both day 3 and 30 (**Fig. S5**). Similarly, the MD data at day 3 showed large portions of the cortex with a small number of rats being significantly different from sham, but this area decreased after harmonization. At day 30, the number of voxels increased around pre-existing areas of difference in the cerebellum and parts of the ipsilateral cortex (**Fig. S10**). In the RD dataset, areas of pre-existing cortical differences decreased after harmonization at day 3 and day 30, except for minor increases in coverage in the contralateral cerebellum (**Fig. S15**).

**Fig. 8.**
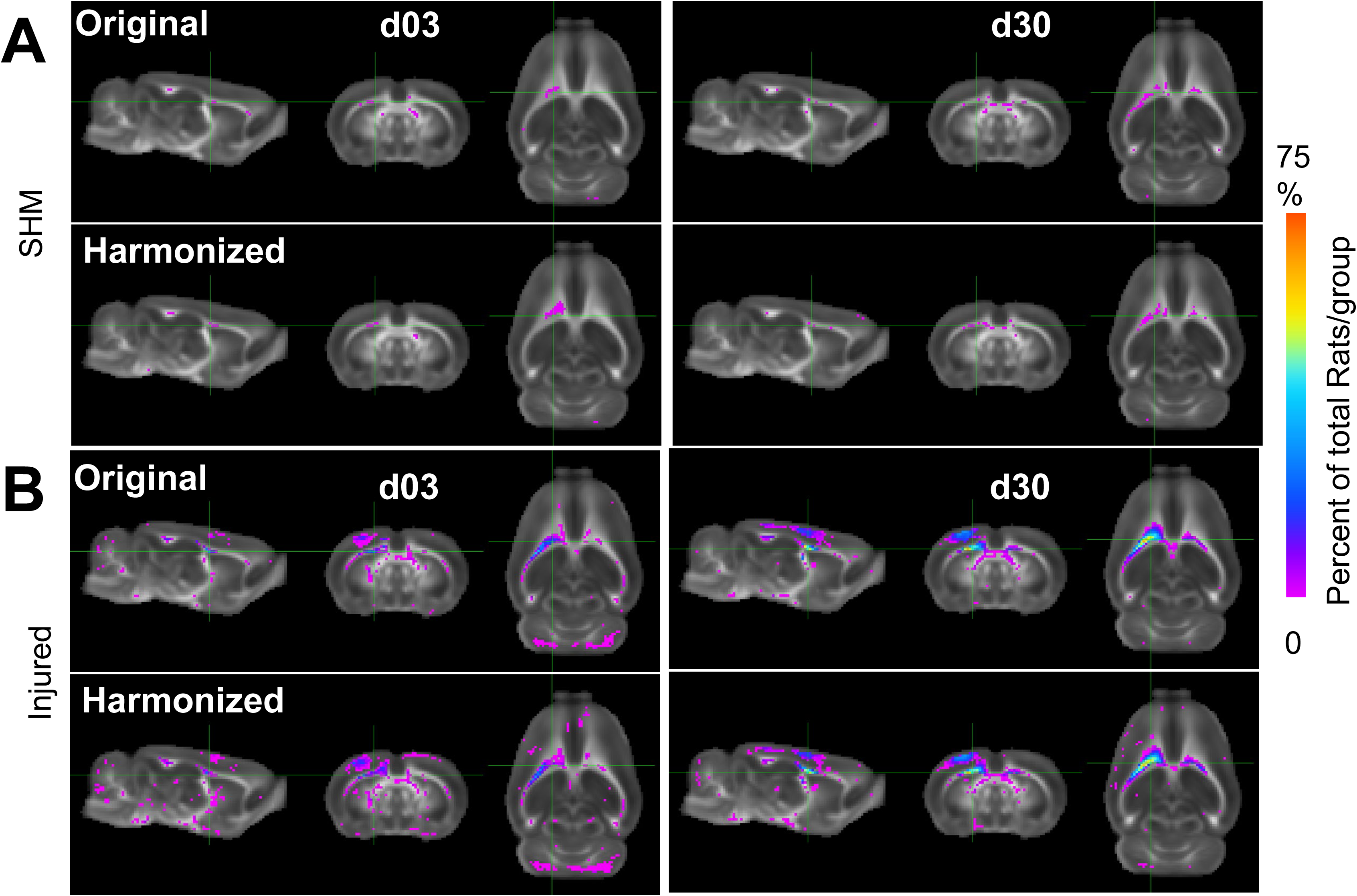
Multi-Site Voxel overlap maps delineating larger regions of common pathology due to data harmonization. **(A)** Sham and **(B)** Injured voxel overlap maps derived from whole-brain data with all three sites showing the incidence of the number of rats at each voxel location where FA values differ significantly from pooled sham data before (label-Original) and after harmonization (label-Harmonized). Images show voxels where there was a lower (Pink) and higher (Blue/Red) proportion of CCI rats in which the FA value was significantly different from pooled shams due to data harmonization (p<0.01, FDR corrected).

## Discussion

This study demonstrates that NeuroCombat harmonization improved the detection of injury after univariate-level harmonization, although this effect was greater using pooled rather than site-specific shams. Despite increases in effect size due to harmonization, group differences were both gained and lost dependent on scalar volume, directional difference compared to sham, and post-injury time. On the other hand, voxel-level harmonization retained group differences present within the original data and led to increases in statistical power and effect size to detect group differences within the primary injured area. These voxel locations displayed a higher frequency of decreased FA in injured rats, but at the potential cost of decreased reliability to detect group differences within regions remote from the injury site.

### Site Outliers Severely Bias NeuroCombat

A significant challenge to NeuroComBat harmonization is the presence of outliers. These findings confirm prior reports (39) that the presence of outliers can severely bias the ability of NeuroCombat to remove site effects. That prior work identified outliers as measurements outside 3 interquartile ranges (IQRs) of Q1 or Q3, and determined that if more than 5% of values at a particular site were outliers, then the ability for NeuroCombat to harmonize data was hindered, and this was associated with reduced variability within the outlier site data (39). Site 4 in the current study exhibited these characteristics with an outlier proportion of >10% when outliers were measured as 1.5 IQRs outside of Q1 and Q3. Difference in scanner field strength is unlikely to be the source of variation since another site that operated at the same field strength (Site 3) showed no such variation. One major difference in hardware between Site 4 and all other sites was the use of different radiofrequency coil hardware that resulted in a reduced brain coverage (**Fig. 1**) that may have accounted for the increased variation. A second major difference was the lower number of rats included from Site 4 in both experimental groups, making the statistical correction to the z-scored volume data larger and thus less sensitive to the detection of abnormal diffusion scalar-related values. This suggests that a more conservative approach during pre-acquisition harmonization is necessary when selecting if site data can be grouped together for harmonization. This may relate to both MRI hardware and injury severity level.

### Choice of Site-Specific versus Pooled Shams is Important

Use of pooled sham data from across all three sites, rather than a site-specific approach, produced a greater percentage of TBI rats where the volume of pathology indicated by FA_Low_ or FA_High_ was greater than in sham rats. This was counter to expectations, where our initial hypothesis was that site-specific sham data would encode site-specific technical, scanner-related effects that would be negated through within-site identification of TBI volumes of pathology, leading to more uniform cross-site data. This premise was based on the current lack of available preclinical phantoms that could be used for correcting for site-specific effects, leaving us to make use of local sham data to fill this gap. However, despite the greater success of using pooled shams, it was only the use of site-specific shams that retained group differences in *univariate-level harmonized* FA_Low_ volumes. Despite this finding, *voxel-level harmonization* reduced cross-site variability, improving injury detection while maintaining all group differences present within the original data. It is possible that the lower sample size at the single-site level could lead to effects that may either overestimate or underestimate true effects, where pooling data can increase sensitivity to true effects (43). It therefore remains crucial to study the effect of how sham data are used when determining the effects of cross-site harmonization.

### Harmonization improves power and effect size in areas with existing biological difference

Consistent with prior literature (12), voxel-level harmonization effectively removed site effects in whole-brain data. As shown in the Bland-Altman plots (Figs. 5B, C), across all combinations of sites and mean FA value of voxel pairs, the difference between each pair of voxels was reduced after harmonization. This reduction in variability across sites after harmonization corresponds with results seen in both the preclinical and clinical spaces (12,17,40). These changes suggest that harmonization improves the ability to detect group differences in areas of pre-existing biological variation. In areas where no significant differences between sham and injury were initially found, power decreased. Univariate measures of pathology showed similar results, with mild effect size increases between injured and sham groups after harmonization. This could explain some of the detrimental effects of harmonization, which has been shown to obscure biological variation in data without site effects, such as in regions unaffected by injury or site (16). The current finding that the proportion of rats which were significantly different from shams did not increase substantially outside of brain regions where group differences were detected before harmonization, confirms that NeuroCombat does not introduce erroneous biological variation (14,15). Voxel-level harmonization appears to only be able to improve detection of group differences across data where group differences already exist, rather than revealing differences obscured by noise or systematic error.

In areas of the brain where group differences were already present, NeuroCombat successfully improved delineation of injury among FA datasets, but not MD, RD or AD. Improvements in power and effect size were seen in areas of the ipsilateral white matter and cortex, which generally corresponds to areas of existing group differences in the FA data. This was further seen by improvements in the proportion of rats that exhibited group differences at these voxels at day 3 and day 30. The use of NeuroCombat on data with site effects may help to improve reproducibility in this respect, as improved power, effect size and more frequent detection of injury can help to reduce erroneous conclusions. An increase in power indicates a benefit for pooling data across sites, allowing for smaller samples from each of multiple sites to confirm observed effects. The decrease in power observed in non-injured areas could have been related to a lack of biological effect in those areas prior to harmonization, where harmonization increased power in areas of existing difference at the expense of whole-brain differences. An alternative reason could be related to a decrease in inaccurate effect sizes within non-injured regions, which would cause power to decrease in those areas and reflect areas of true biological difference. Similar findings have been reported in clinical FA data: harmonization with NeuroCombat was able to increase the number of voxels associated with age in a clinical multi-site and multi-scanner study of DTI measures’ relationship with age (12), as well as improve power in the detection of differences among individuals with schizophrenia in T1 derived metrics of cortical thickness, surface area and subcortical volumes (5). The present study builds upon these initial findings by validating NeuroCombat’s efficacy in preclinical data, as well as by demonstrating that these improvements in the detection of group differences are confined within areas of existing biological variation.

There was a relative lack of success by harmonization to improve injury detection among AD, MD and RD datasets compared to FA, despite varying degrees of power and effect size improvements. Reasons for this are not clear, but it may relate to site differences in measurement of absolute diffusivities compared to a more equitable comparison of FA which is a normalized ratio of these diffusivities, and thus less likely to differ in absolute values across site. Further examination into whether normalization of unbounded metrics helps to improve the efficacy of NeuroCombat may be warranted to better understand this phenomenon. Other methods of ComBat that have been used in imaging data include generalized additive models (gamCombat) which accounts for non-linear covariates, longCombat which can be used to detect changes over time, and gaussian mixture model-combat that removes variation due to unidentified covariates (16,41). NeuroCombat has been shown to be preferable when conducting cross-sectional data harmonization compared to longCombat (16). While in the present study we were focused on group-level comparisons at each timepoint, the use of longCombat may well be useful to capture differences in injury trajectories across site in future work.

## Limitations

Inherent in the preprocessing of image data are the potential errors associated with the requirement to conduct spatial transformations to ensure correct registration to a common 3D space. Multiple transformations could have impacted the quality and precision of the resulting measures of pathology. The heterogeneous nature of injury may also play a role in the decrease in power in non-significantly different brain areas. Due to potential minor differences in delivery of trauma, brain areas of some rats may have been more injured compared to similar areas in other rats. This may lead to less consistent measurements and interfere with NeuroCombat’s abilities to resolve site effects.

Univariate level volume analysis was conducted after identifying regions of Scalar_Low_ and Scalar_High_ through a statistical-based z-corrected threshold (29, 42). However, there is no agreed upon statistical correction required for the analysis of voxel-based data. It is possible that the strict threshold used here (P<0.001) reduced the ability to accurately judge pathologic tissue within injured versus sham rats, as indicated by the low overlap between rats in site 4 (**Fig.1**). More work is required to determine the impact of statistical correction threshold on determining group differences after TBI.

The biological basis for FA_HIGH_ regions after TBI is uncertain, although it has been found before in the same spatially discrete patterns from FA_LOW_ regions (44). Noteworthy here though is that the benefit provided by multi-site harmonization was afforded by FA_LOW_, and from which the biological underpinnings is far more certain.

## Conclusion

NeuroCombat harmonization demonstrates utility in reducing site effects in rodent imaging across multiple sites and scanner field strengths. By increasing the statistical power and effect size to detect areas of injury, post-acquisition, cross-site data harmonization improves the ability to discriminate between sham and injured rats. This should enable improved efficiency for preclinical study completion by collecting data at multiple sites.

## Author Contribution Statement

GK, RF, NGH: Methodology, Formal Analysis, Visualization, Writing.

AVK, JZ, MF, RCK, MPB, JM, AP, TA, BM, AB, JW, AP, YZ, NGH: Data acquisition and/or curation

AVK, JZ, MF, RCK, MPB, JM, ARF, IBW, NGH: Review and Editing.

NGH: Conceptualization, project administration.

NGH, IBW, JZ, RCK, MPB, JM, KKW, MF, ARF, JRH: funding.

## Acknowledgments

The TOP-NT Investigators include (in alphabetical order):

Albenese, Chris, Georgetown University, Washington, DC Buenaventura, Ruchelle, Georgetown University, Washington,

Dickstein, Dara L., Uniformed Services University of the Health Sciences, Bethesda, MD

Fan, Anya, Uniformed Services University of the Health Sciences and the Henry M. Jackson Foundation for the Advancement of Military Medicine, Bethesda, MD

Fu, Amanda, Uniformed Services University of the Health Sciences, Bethesda, MD and the Henry M. Jackson Foundation for the Advancement of Military Medicine Harvey, Alexander C., Georgetown University, Washington, DC

Labastida, Javier Allende, Johns Hopkins University, Baltimore, Maryland Lee, Yichien, Georgetown University, Washington, DC

Liu, Jiong, Uniformed Services University of the Health Sciences and the Henry M. Jackson Foundation for the Advancement of Military Medicine, Bethesda, MD

Klee, Madison P., University of California Los Angeles, Los Angeles, CA Main, Bevan S., Georgetown University, Washington, DC

Rodruigez, Olga, Georgetown University, Washington, DC

Tong, Jonathan, University of California Los Angeles, Los Angeles, CA Torres-Espín, Abel, University of Waterloo, Waterloo, ON

Tucker, Laura B., Uniformed Services University of the Health Sciences and the Henry M. Jackson Foundation for the Advancement of Military Medicine, Bethesda, MD

Vanderveer-Harris, Nathan, University of California Los Angeles, Los Angeles, CA

## Supplementary Figures Legend

**Fig. S1.**
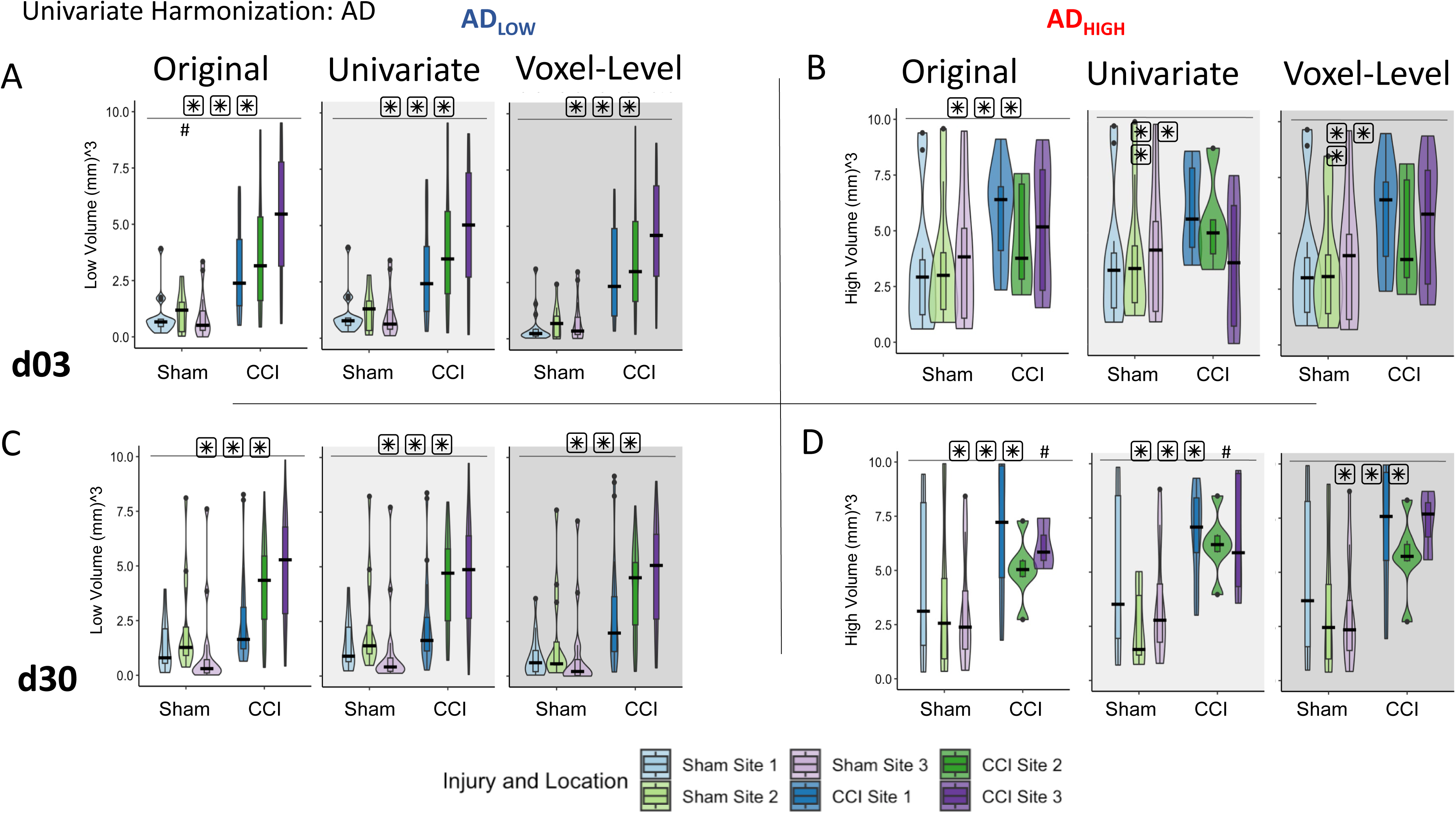
Effect of univariate and voxel-level harmonization on the burden of whole brain injury volume using pooled sham data: AD. Brain volumes of pathology identified by (**A,C**) AD_Low_ and (**B,D**) AD_High_ at 3d (**A,B**) and 30d (**C,D**) are plotted for each site before (“Original”) and after univariate and voxel-level harmonization. ###/*** = a significant effect of location and injury, respectively, p<0.001, ##/** = p<0.01, #/* = p<0.05 (linear mixed model ANOVA).

**Fig S2.**
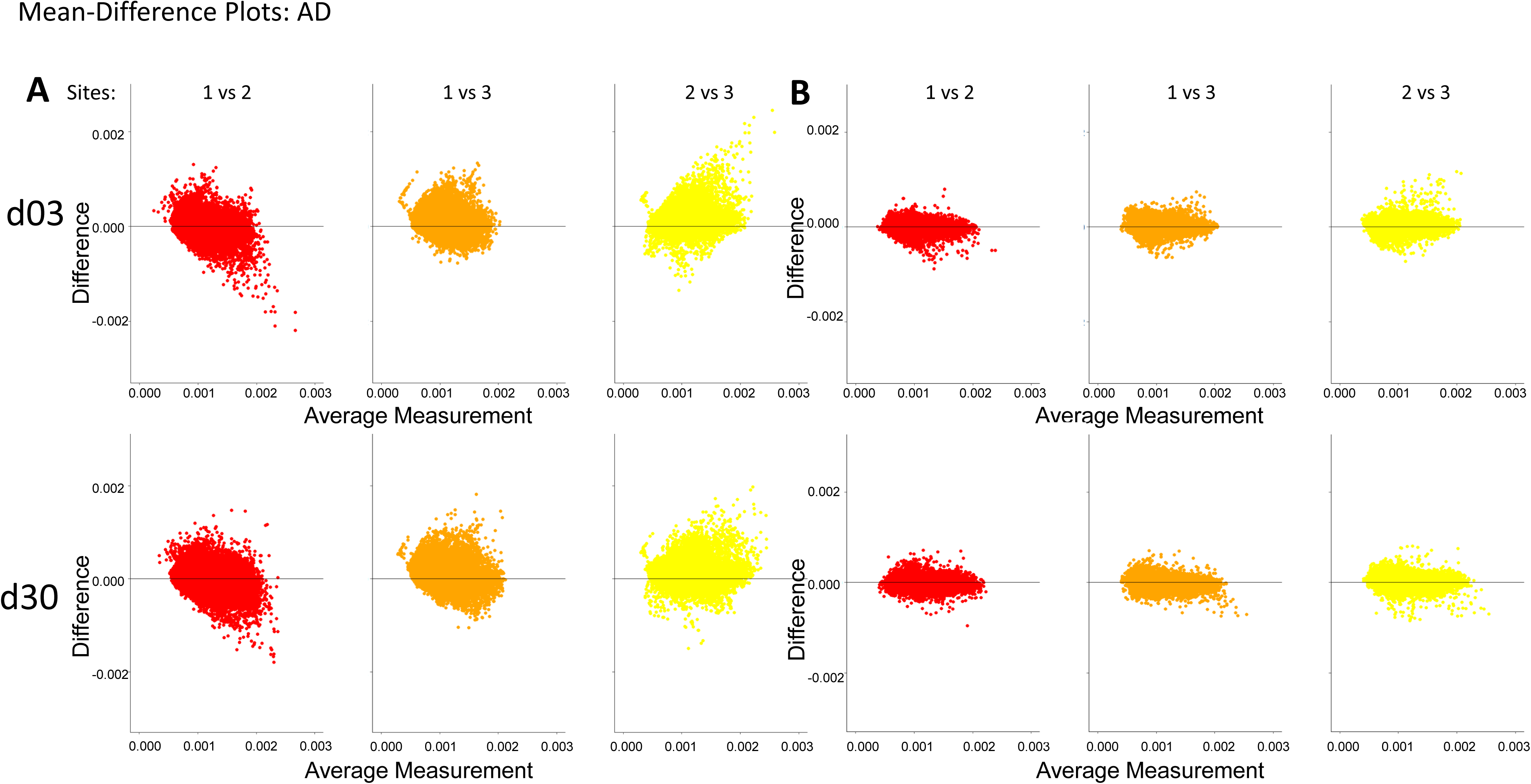
Harmonization at voxel-level resolution: AD. **(A-B**) Harmonization of AD values of each individual voxel across site pairs is demonstrated using Bland-Altman plots for (**A**)original, unharmonized values and (**B**) harmonized values. The difference between sites decreases due to harmonization.

**Fig S3.**
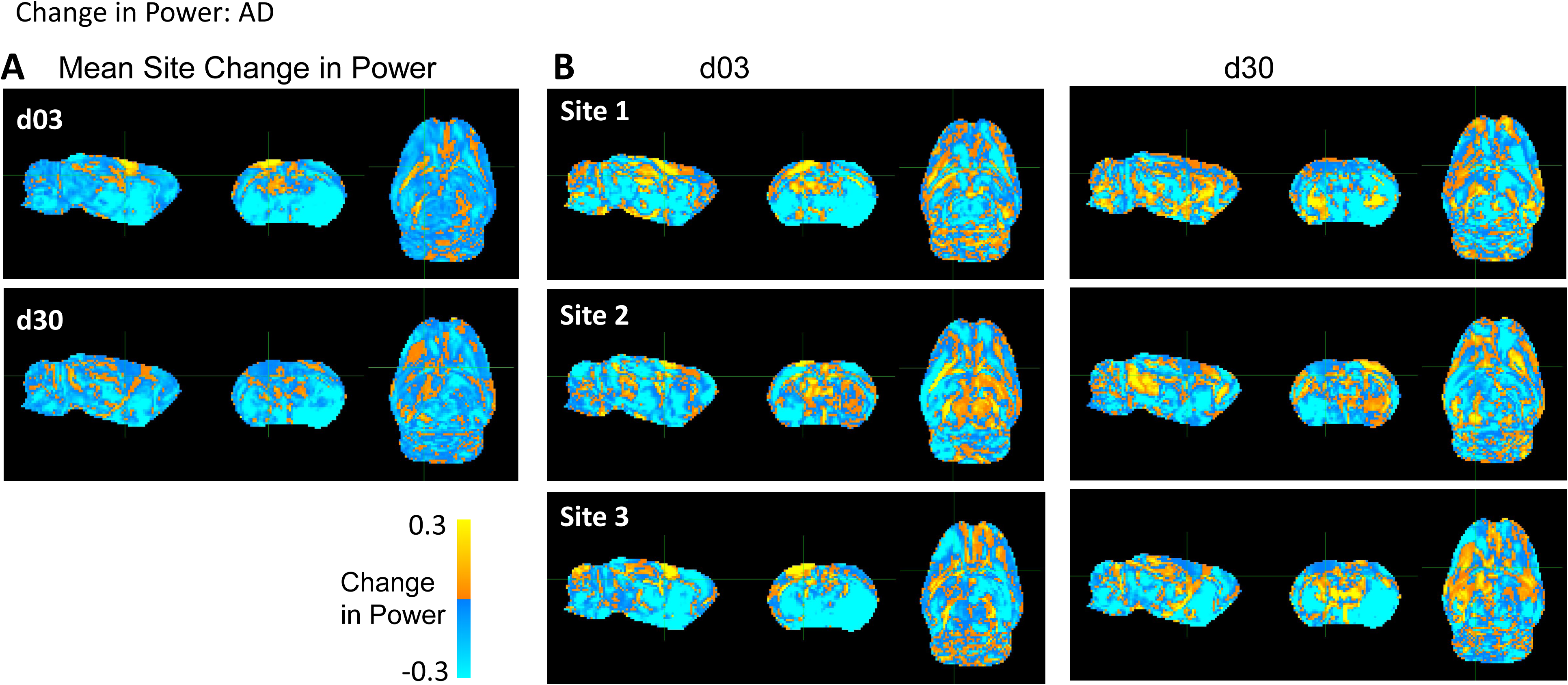
Whole-brain maps of regional changes in power due to harmonization: AD. **(A)** Difference in power across a pooled average of all three sites post-injury due to harmonization and **(B)** across site and post-injury day. The magnitude scale -30% and 30% indicates the percent difference in statistical power between pre and post voxel-level harmonization, where yellow/blue indicate power increases and decreases, respectively due to harmonization.

**Fig. S4.**
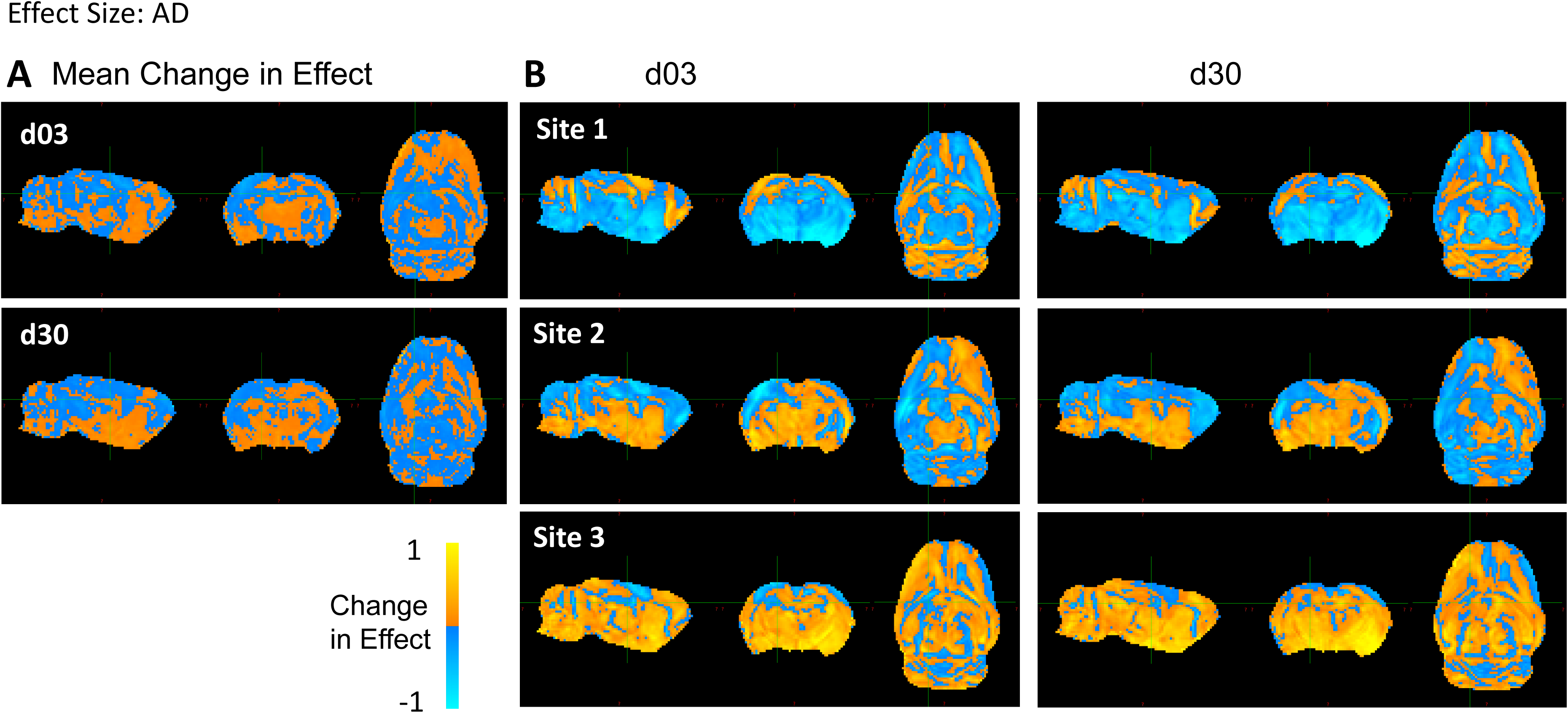
Whole-brain maps of regional changes in effect size due to harmonization: AD. **(A)** Difference in effect size across a pooled average of all three site post-injury due to harmonization and **(B)** across site and post-injury day. The magnitude scale –1 and +1 Cohen’s d indicates the difference in effect size/voxel between pre and post voxel-level harmonization, where yellow/blue indicate power increases and decreases, respectively due to harmonization.

**Fig. S5.**
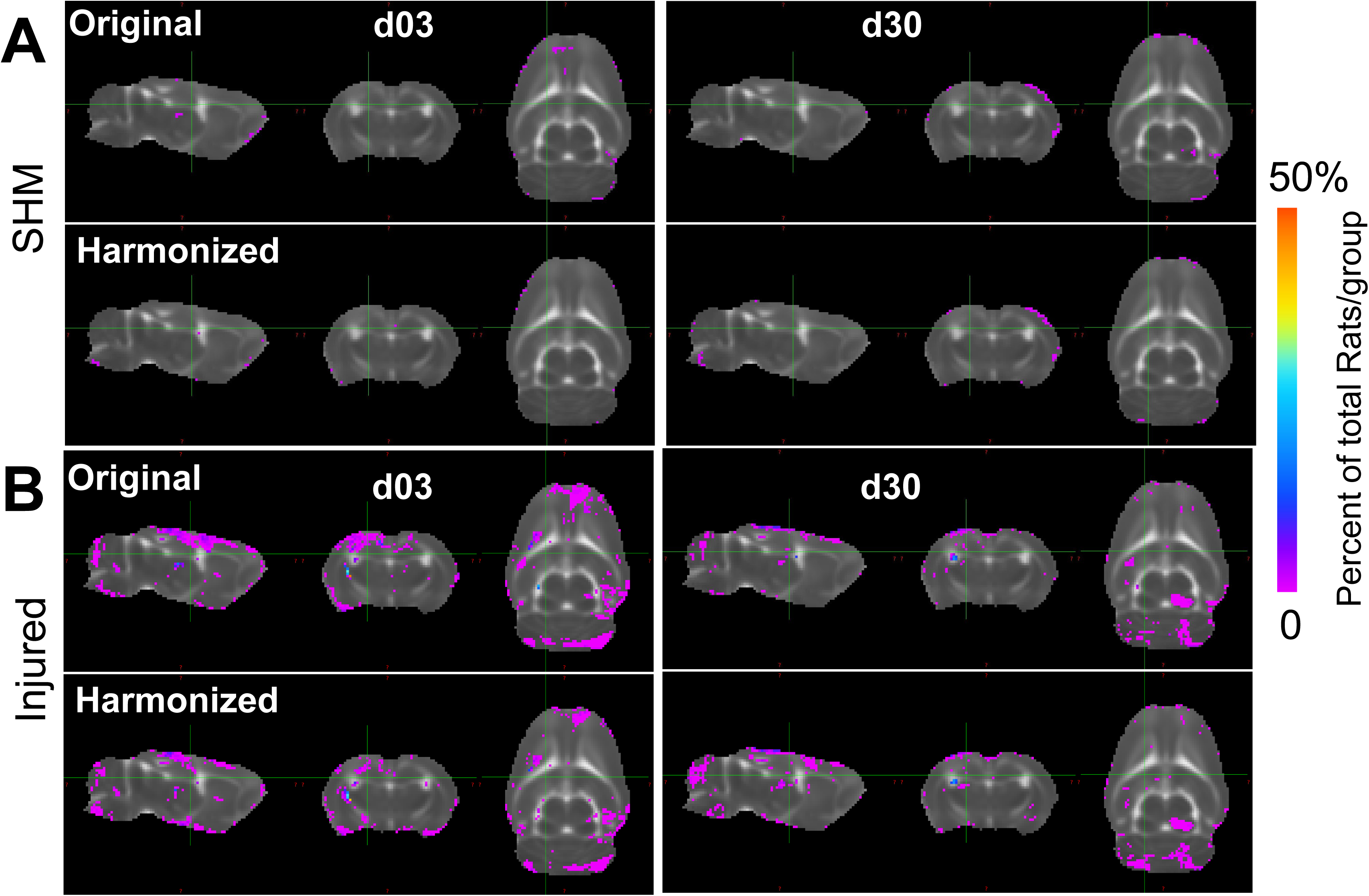
Multi-Site Voxel overlap maps delineating regions of common pathology due to data harmonization: AD. Voxel overlap maps derived from whole-brain data across all 3 sites showing the incidence of the number of rats at each voxel location where AD values differ significantly from pooled sham data **(A)** before (Original) and **(B)** after harmonization (Harmonized). Images show voxels where there was a lower (Pink) and higher (Blue/Red) proportion of rats in which the AD value was significantly different from pooled shams due to data harmonization (p<0.01, FDR corrected).

**Fig. S6.**
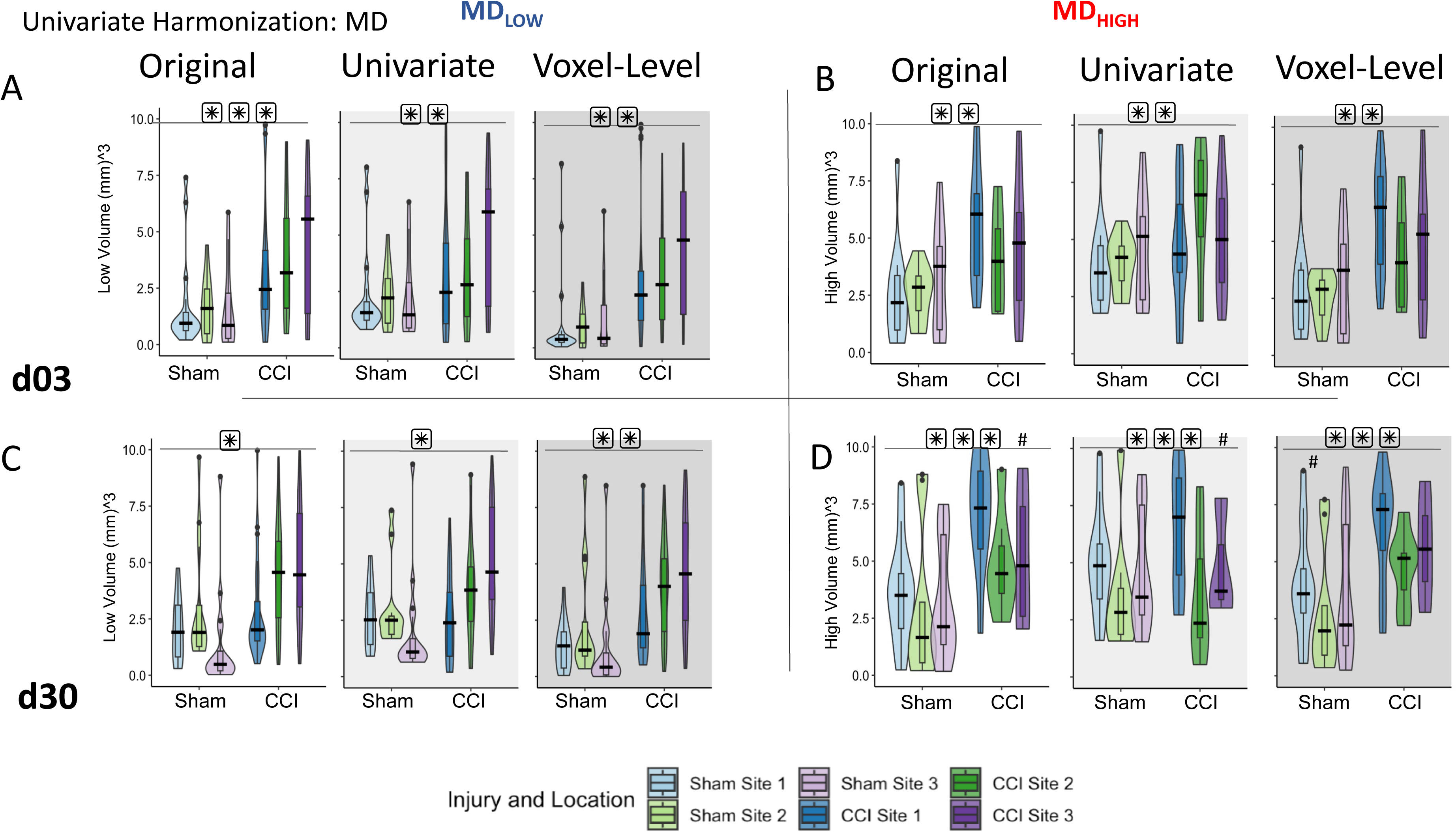
Effect of univariate and voxel-level harmonization on the burden of whole brain injury volume using pooled sham data: MD. Brain volumes of pathology identified by (**A,C**) MD_Low_ and (**B,D**) MD _High_ at 3d (**A,B**) and 30d (**C,D**) are plotted for each site before (“Original”) and after univariate and voxel-level harmonization. ###/*** = a significant effect of location and injury, respectively, p<0.001, ##/** = p<0.01, #/* = p<0.05 (linear mixed model ANOVA).

**Fig S7.**
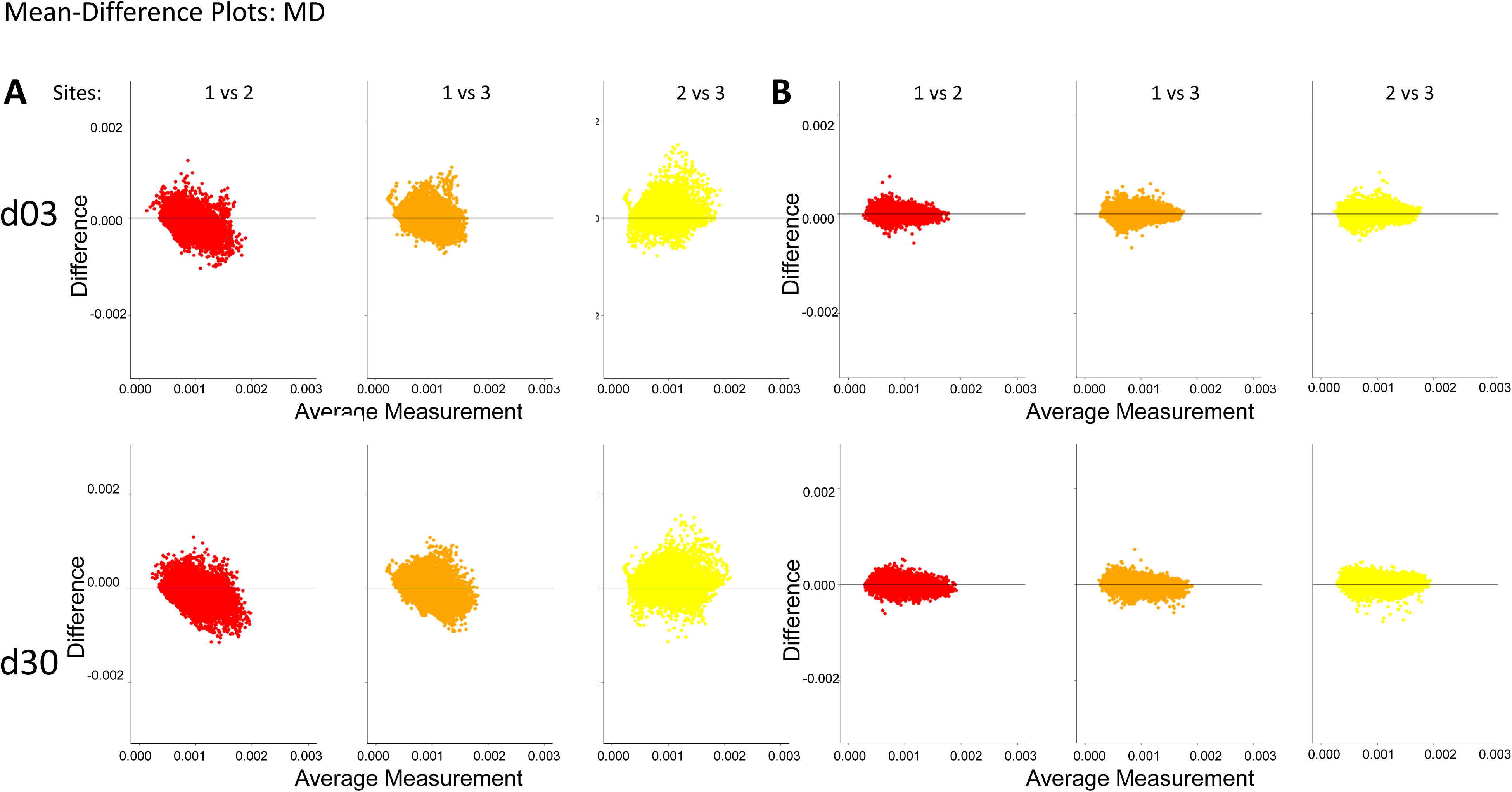
Harmonization at voxel-level resolution: MD. **(A-B**) Harmonization of MD values of each individual voxel across site pairs is demonstrated using Bland-Altman plots for (**A**)original, unharmonized values and (**B**) harmonized values. The difference between sites decreases due to harmonization.

**Fig S8.**
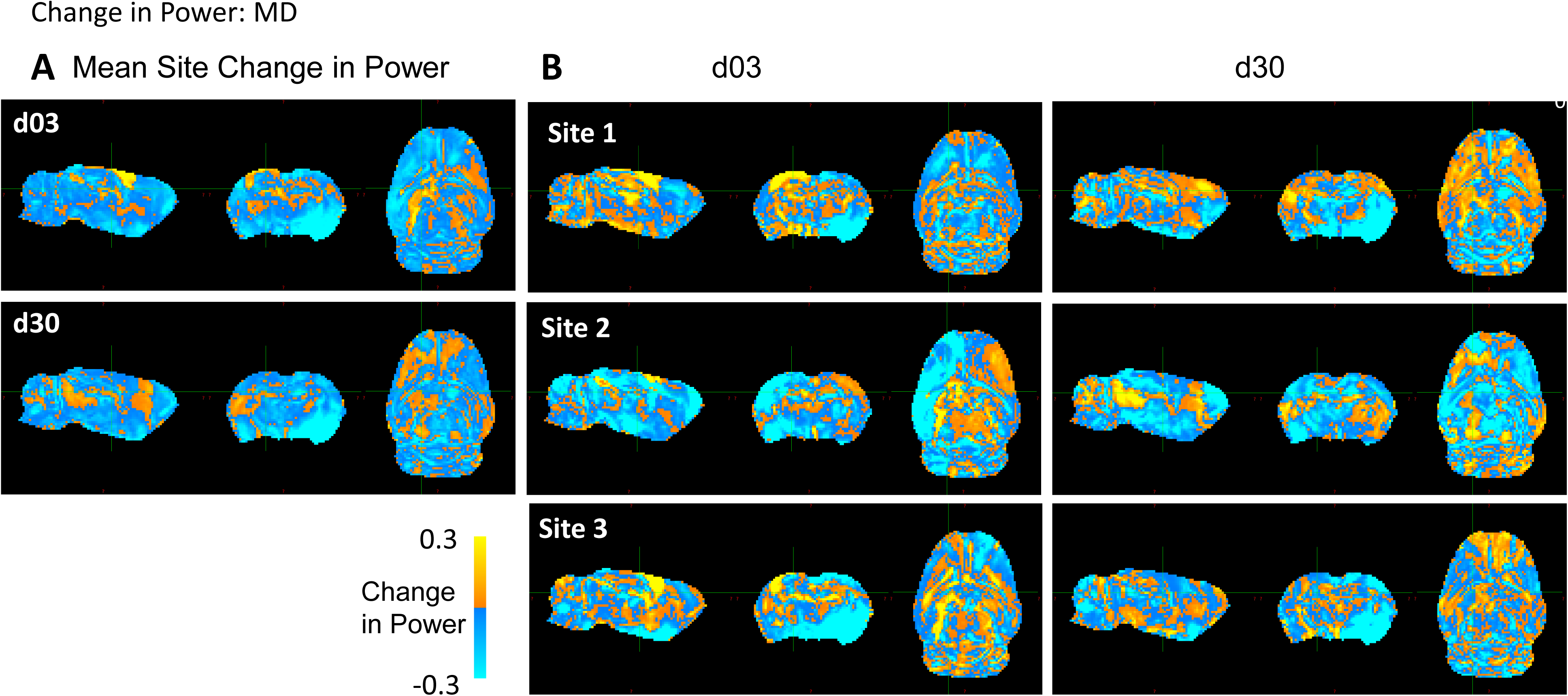
Whole-brain maps of regional changes in power due to harmonization: MD. **(A)** Difference in power across a pooled average of all three sites post-injury due to harmonization and **(B)** across site and post-injury day. The magnitude scale -30% and 30% indicates the percent difference in statistical power between pre and post voxel-level harmonization, where yellow/blue indicate power increases and decreases, respectively due to harmonization.

**Fig. S9.**
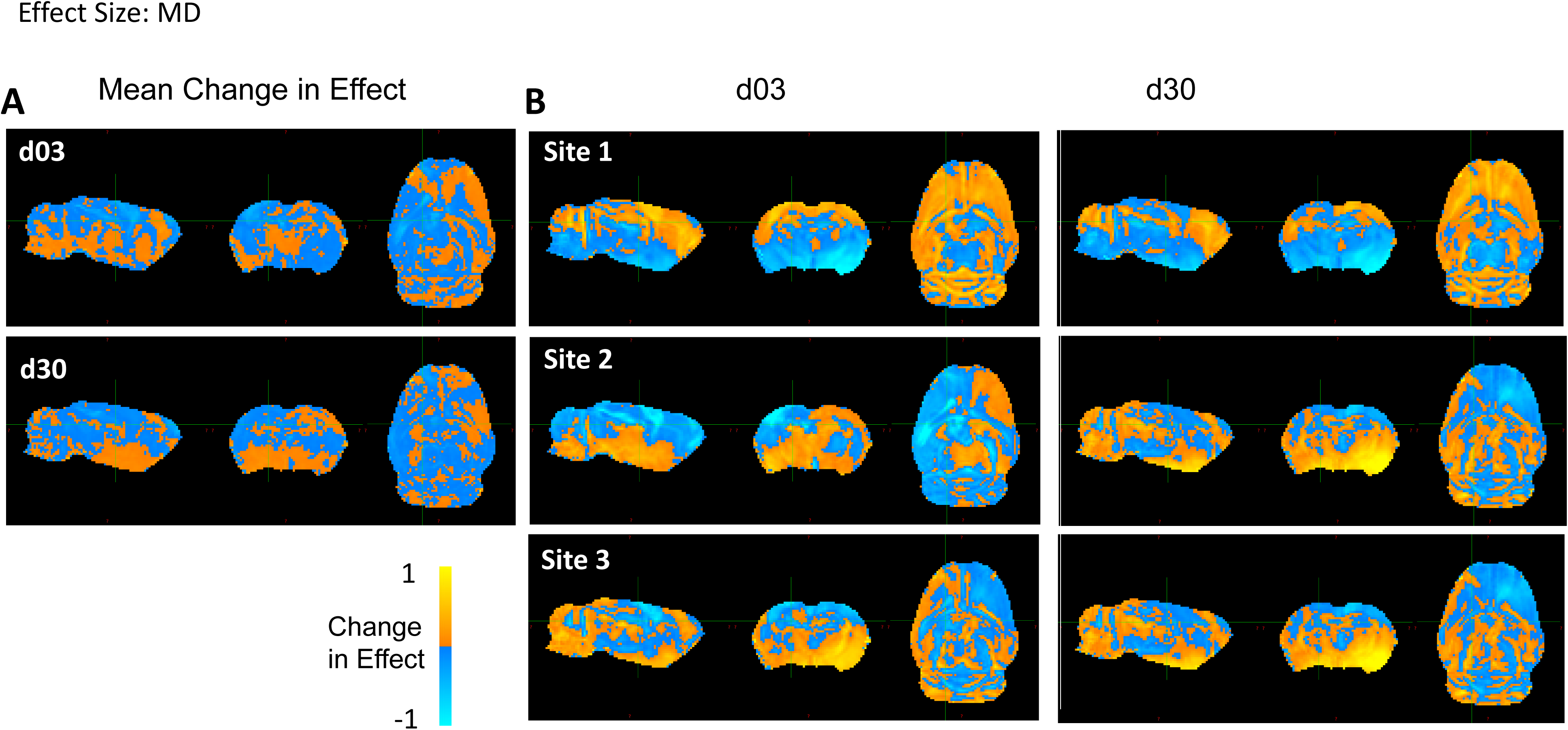
Whole-brain maps of regional changes in effect size due to harmonization: MD. **(A)** Difference in effect size across a pooled average of all three site post-injury due to harmonization and **(B)** across site and post-injury day. The magnitude scale –1 and +1 Cohen’s d indicates the difference in effect size/voxel between pre and post voxel-level harmonization, where yellow/blue indicate power increases and decreases, respectively due to harmonization.

**Fig. S10.**
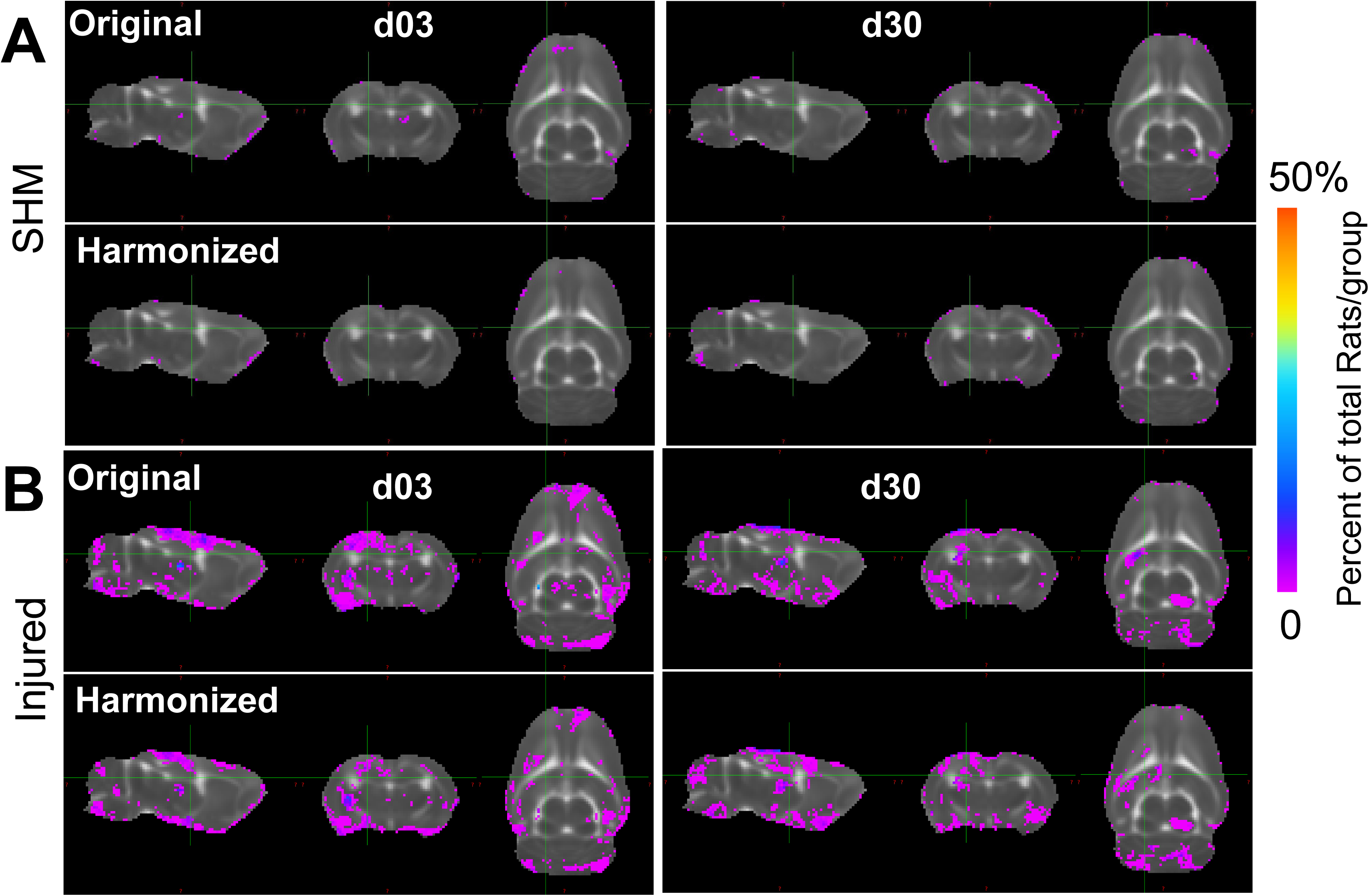
Multi-Site Voxel overlap maps delineating regions of common pathology due to data harmonization: MD. Voxel overlap maps derived from whole-brain data across all 3 sites showing the incidence of the number of rats at each voxel location where MD values differ significantly from pooled sham data **(A)** before (Original) and **(B)** after harmonization (Harmonized). Images show voxels where there was a lower (Pink) and higher (Blue/Red) proportion of rats in which the MD value was significantly different from pooled shams due to data harmonization (p<0.01, FDR corrected).

**Fig. S11.**
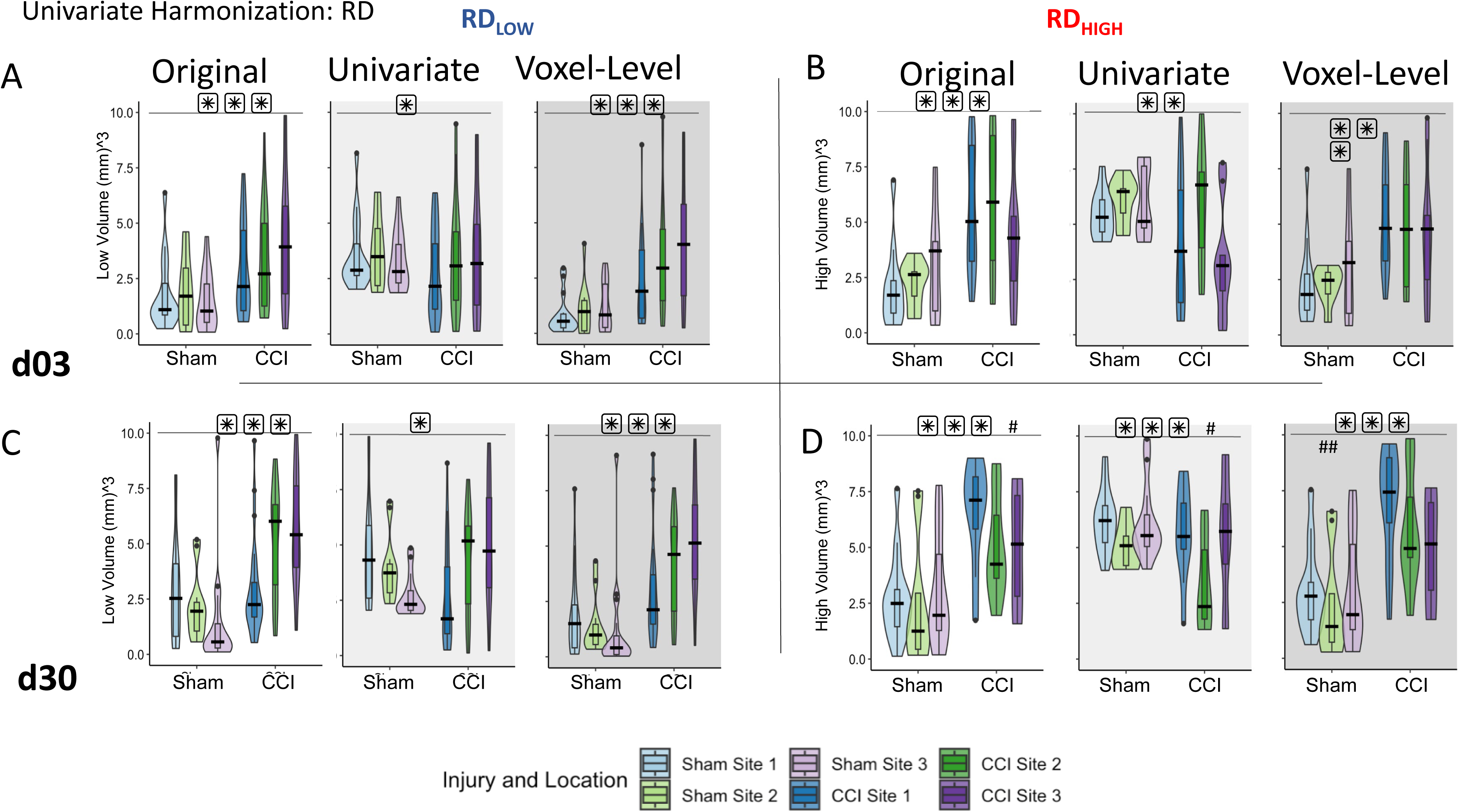
Effect of univariate and voxel-level harmonization on the burden of whole brain injury volume using pooled sham data: RD. Brain volumes of pathology identified by (**A,C**) RD_Low_ and (**B,D**) RD_High_ at 3d (**A,B**) and 30d (**C,D**) are plotted for each site before (“Original”) and after univariate and voxel-level harmonization. ###/*** = a significant effect of location and injury, respectively, p<0.001, ##/** = p<0.01, #/* = p<0.05 (linear mixed model ANOVA).

**Fig S12.**
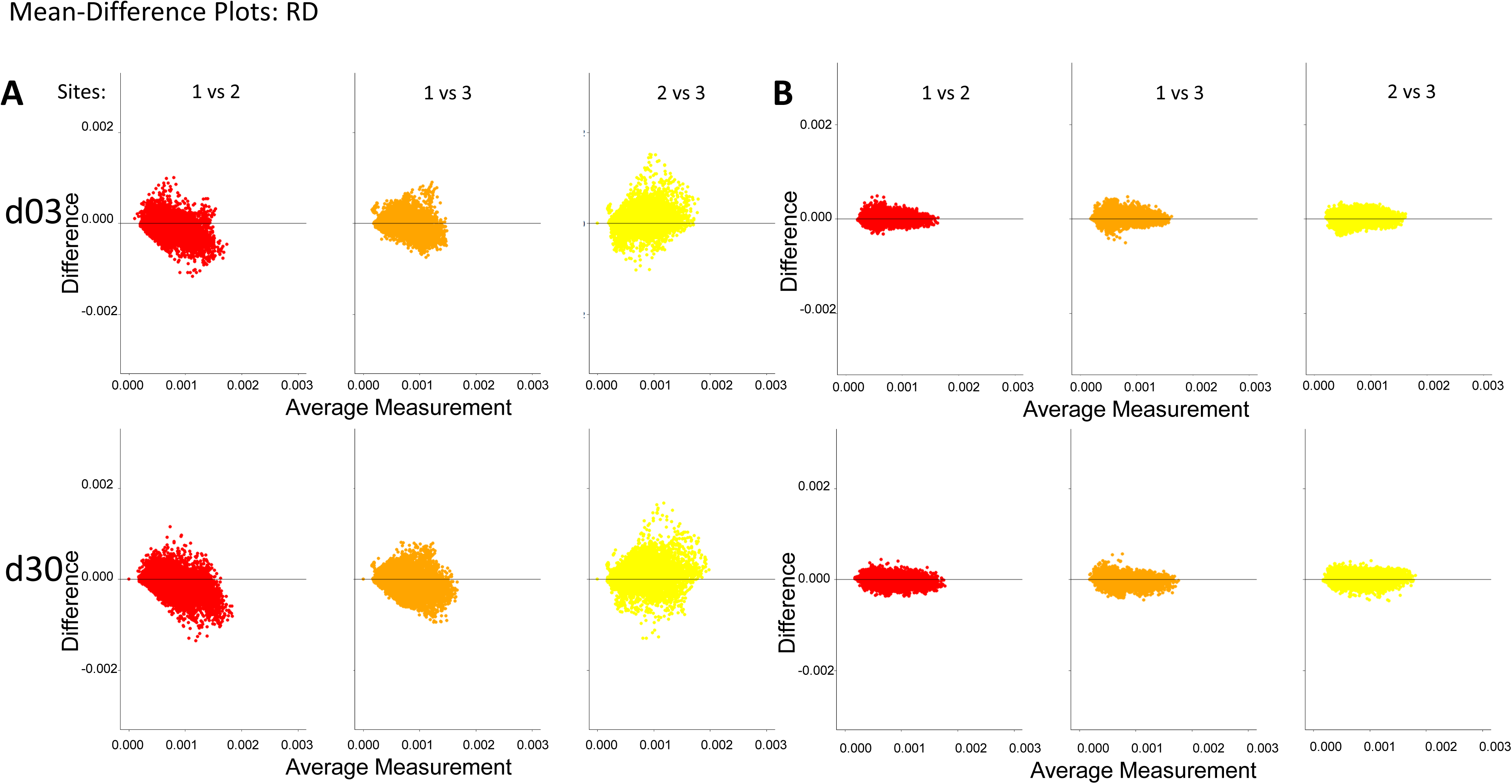
Harmonization at voxel-level resolution: RD. **(A-B**) Harmonization of RD values of each individual voxel across site pairs is demonstrated using Bland-Altman plots for (**A**)original, unharmonized values and (**B**) harmonized values. The difference between sites decreases due to harmonization.

**Fig S13.**
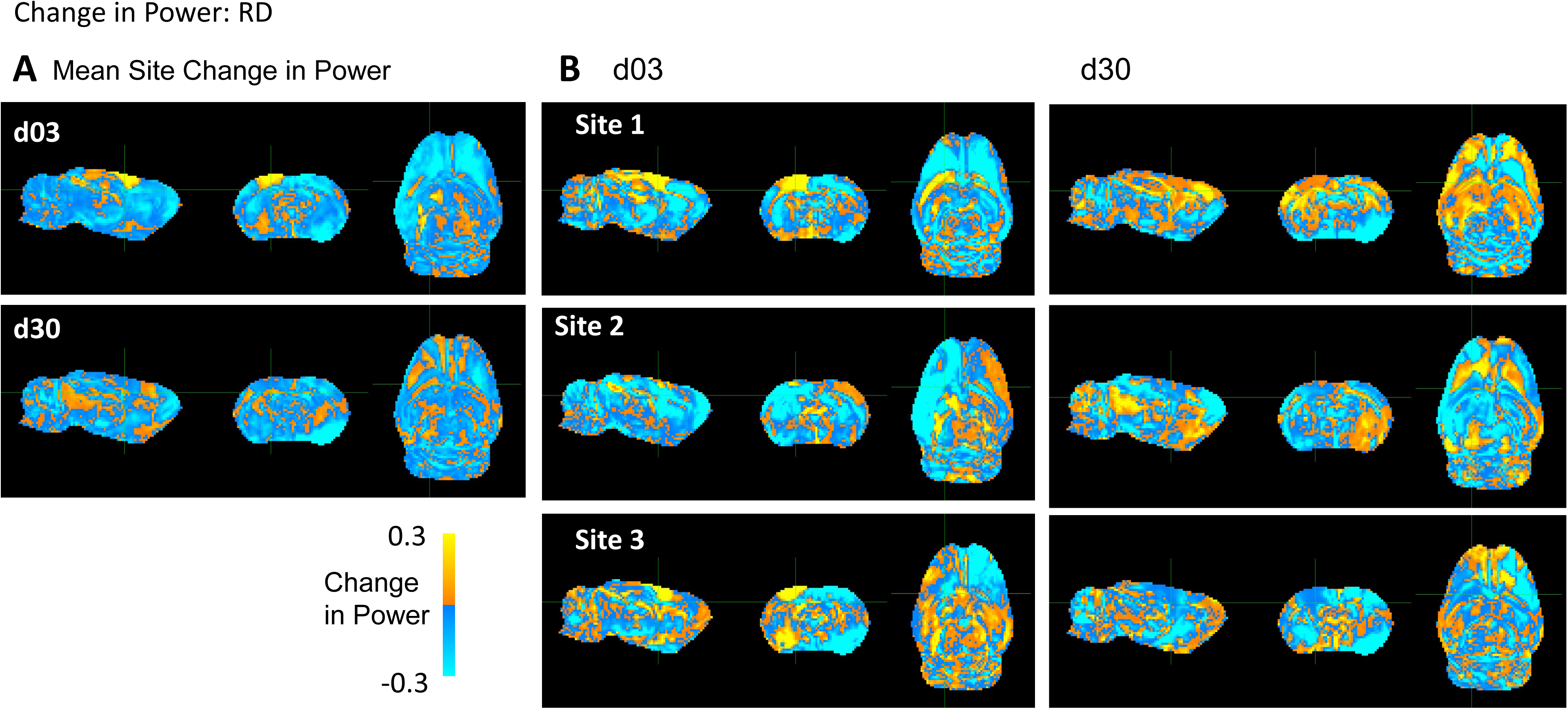
Whole-brain maps of regional changes in power due to harmonization: RD. **(A)** Difference in power across a pooled average of all three sites post-injury due to harmonization and **(B)** across site and post-injury day. The magnitude scale -30% and 30% indicates the percent difference in statistical power between pre and post voxel-level harmonization, where yellow/blue indicate power increases and decreases, respectively due to harmonization.

**Fig. S14.**
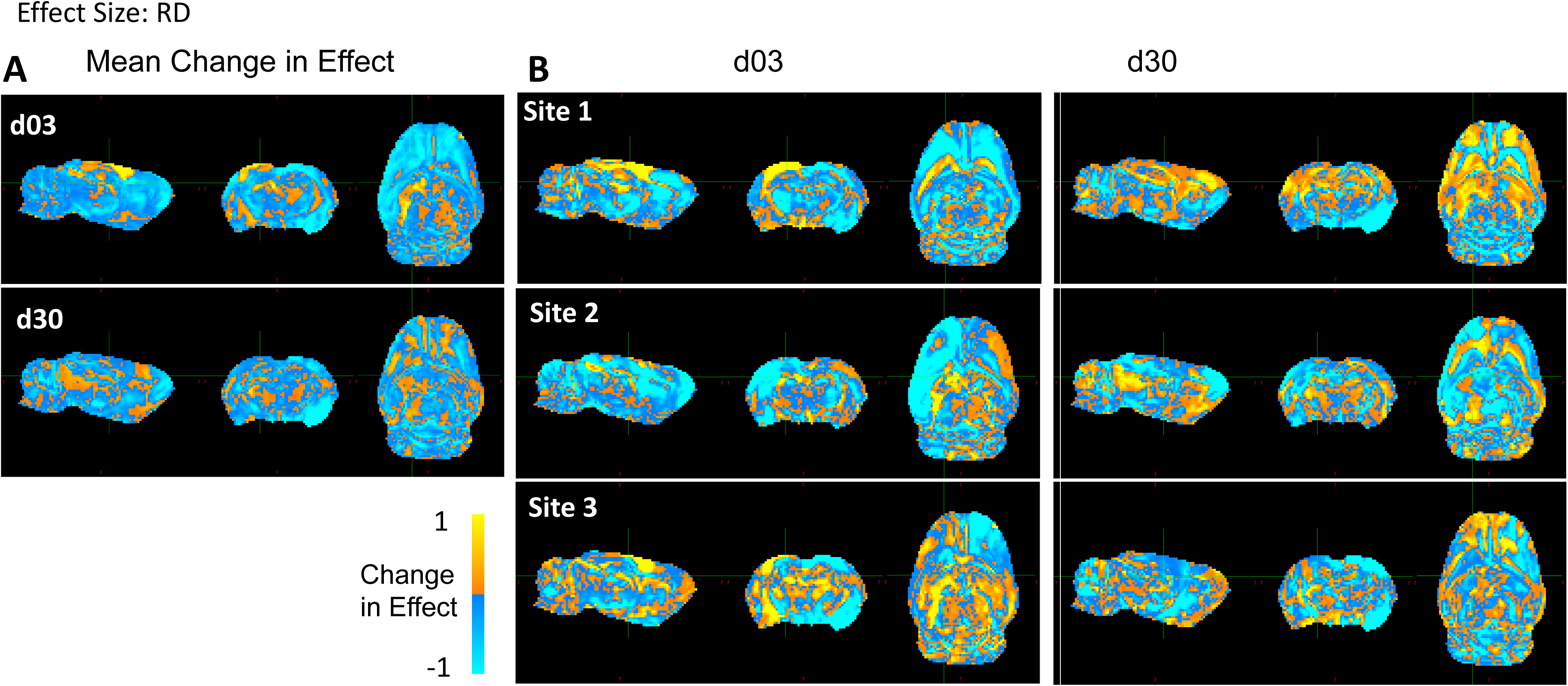
Whole-brain maps of regional changes in effect size due to harmonization: RD. **(A)** Difference in effect size across a pooled average of all three site post-injury due to harmonization and **(B)** across site and post-injury day. The magnitude scale –1 and +1 Cohen’s d indicates the difference in effect size/voxel between pre and post voxel-level harmonization, where yellow/blue indicate power increases and decreases, respectively due to harmonization.

**Fig. S15.**
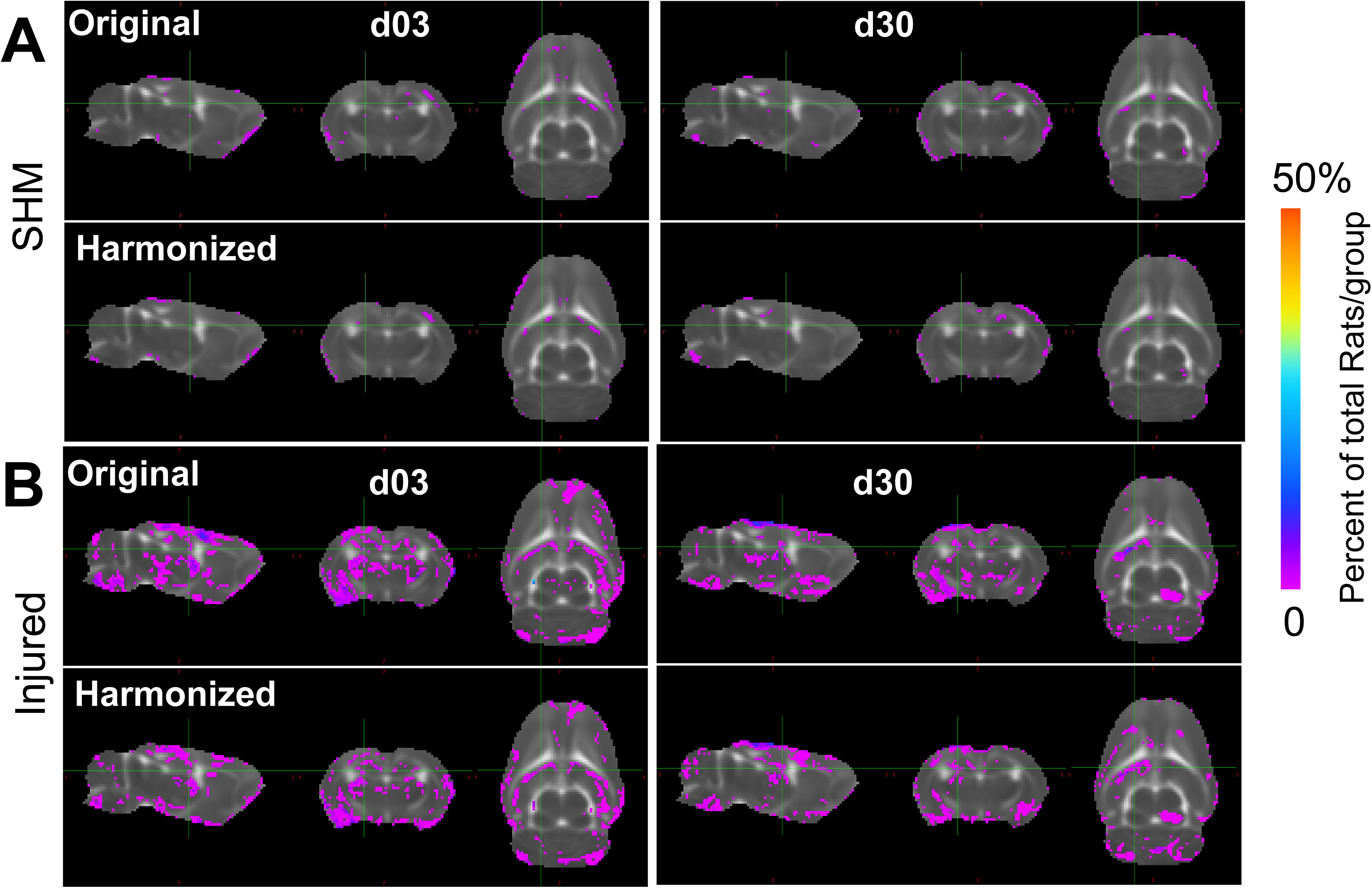
Multi-Site Voxel overlap maps delineating regions of common pathology due to data harmonization: RD. Voxel overlap maps derived from whole-brain data across all 3 sites showing the incidence of the number of rats at each voxel location where RD values differ significantly from pooled sham data **(A)** before (Original) and **(B)** after harmonization (Harmonized). Images show voxels where there was a lower (Pink) and higher (Blue/Red) proportion of rats in which the RD value was significantly different from pooled shams due to data harmonization (p<0.01, FDR corrected).

**Fig. S16.**
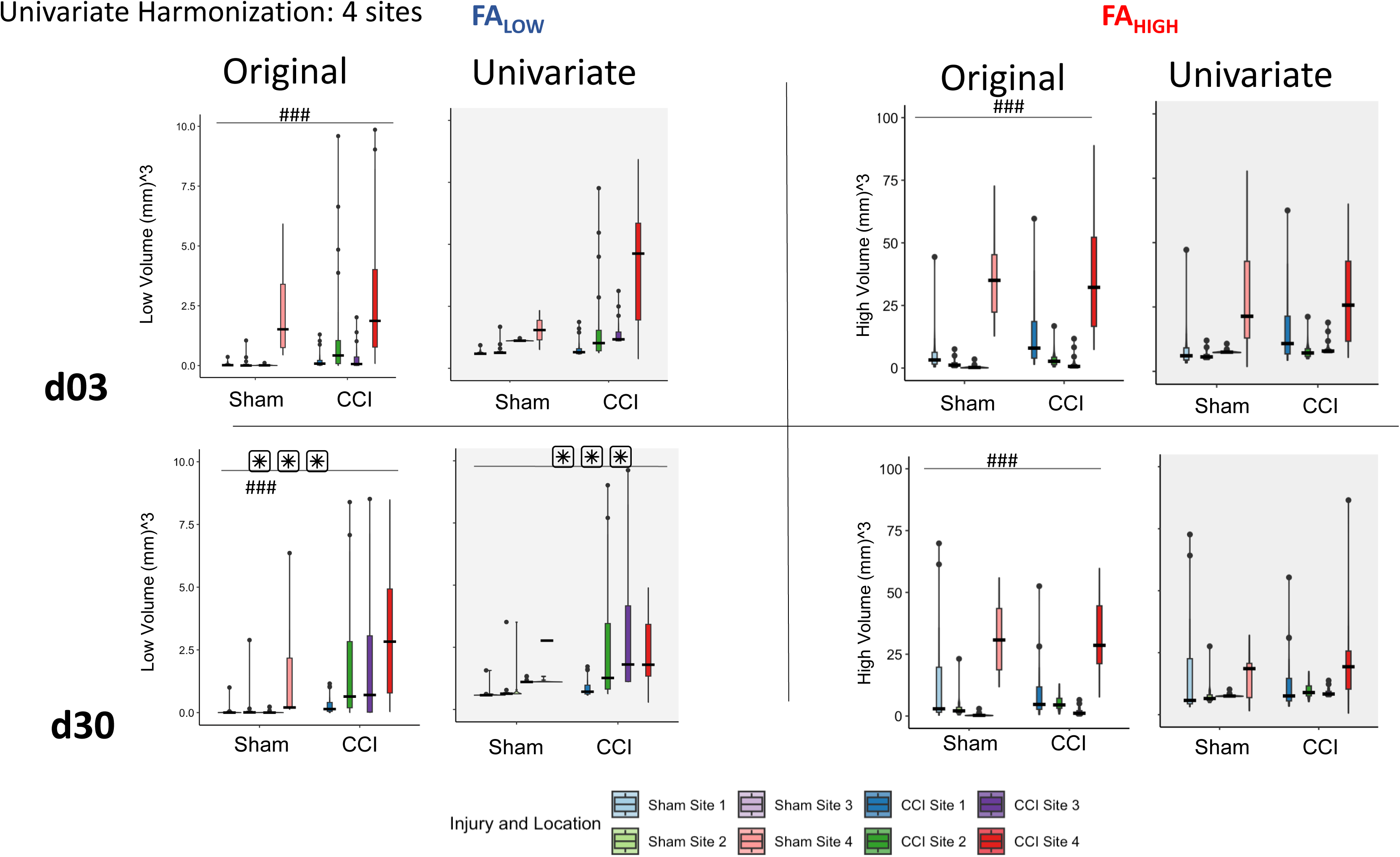
Effect of univariate harmonization on the burden of whole brain injury volume using pooled sham data at all four sites. Brain volumes of pathology at all four sites identified by (**A,C**) FA_Low_ and (**B,D**) FA_High_ at 3d (**A,B**) and 30d (**C,D**) are plotted for each site before (“Original”) and after univariate and voxel-level harmonization. ###/*** = a significant effect of location and injury, respectively, p<0.001, ##/** = p<0.01, #/* = p<0.05 (linear mixed model ANOVA).

**Fig. S17.**
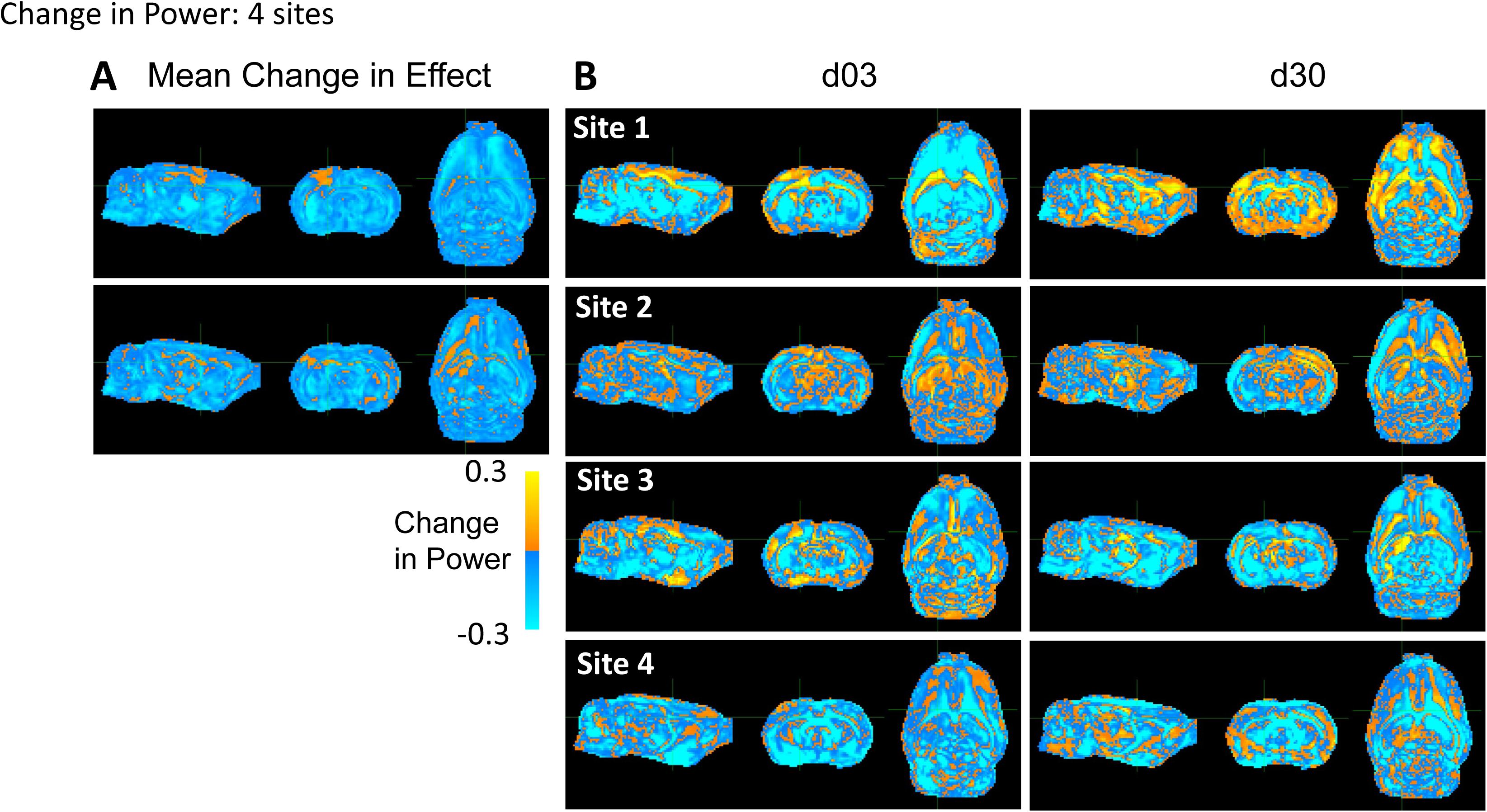
Whole-brain maps of regional changes in effect size due to harmonization across all four sites. **(A)** Difference in effect size across a pooled average of all four site post-injury due to harmonization and **(B)** across site and post-injury day. The magnitude scale –1 and +1 Cohen’s d indicates the difference in effect size/voxel between pre and post voxel-level harmonization, where yellow/blue indicate power increases and decreases, respectively due to harmonization.

**Fig. S18.**
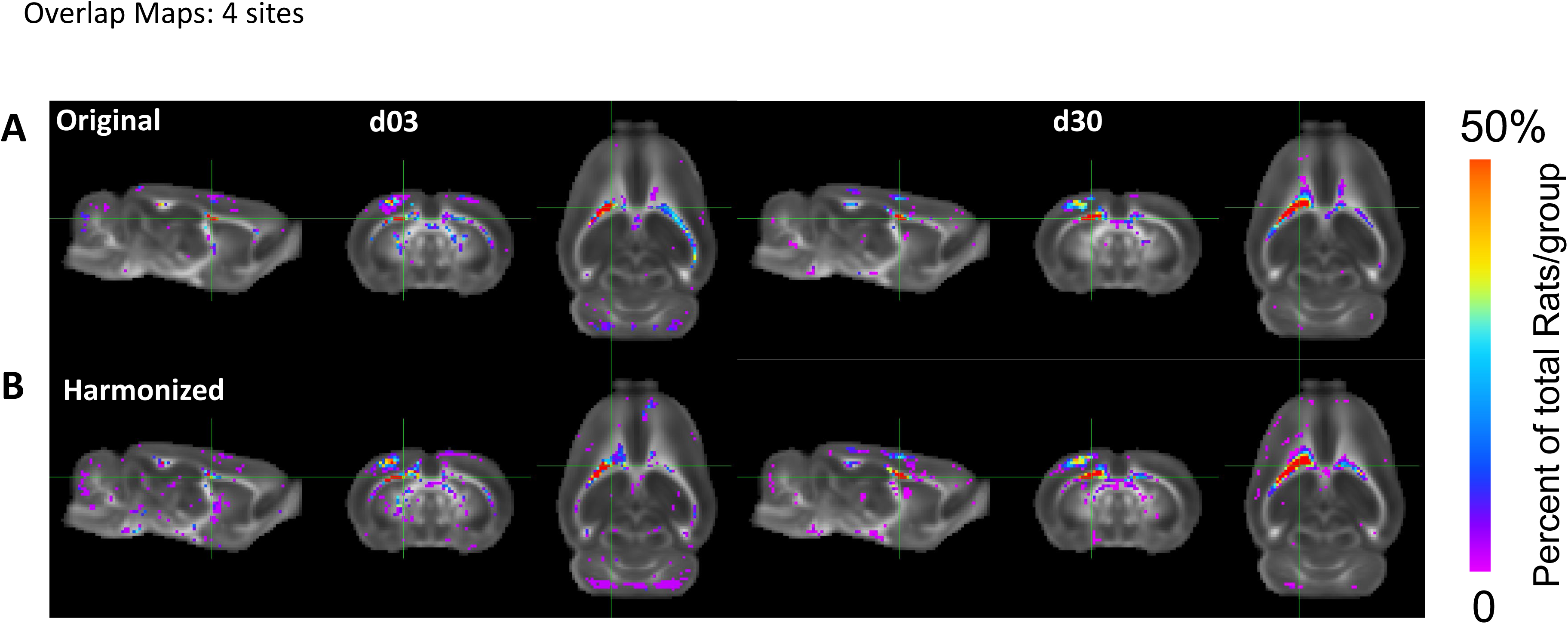
Multi-Site Voxel overlap maps delineating regions of common pathology due to data harmonization at all four sites. Voxel overlap maps derived from whole-brain data across all four sites showing the incidence of the number of rats at each voxel location where FA values differ significantly from pooled data **(A)** before (Original) and **(B)** after harmonization (Harmonized). Images show voxels where there was a lower (Pink) and higher (Blue/Red) proportion of rats in which the FA value was significantly different from pooled shams due to data harmonization (p<0.01, FDR corrected).

## References

1. Albensi BC, Knoblach SM, Chew BGM, O’Reilly MP, Faden AI, Pekar JJ. Diffusion and High Resolution MRI of Traumatic Brain Injury in Rats: Time Course and Correlation with Histology. Exp Neurol. 2000 Mar 1;162(1):61–72.

2. Benson RR, Meda SA, Vasudevan S, Kou Z, Govindarajan KA, Hanks RA, et al. Global white matter analysis of diffusion tensor images is predictive of injury severity in traumatic brain injury. J Neurotrauma. 2007 Mar;24(3):446–59.

3. Harris NG, Verley DR, Gutman BA, Sutton RL. Bi-directional changes in fractional anisotropy after experiment TBI: Disorganization and reorganization? NeuroImage. 2016 Jun;133:129–43.

4. Basser PJ, Pierpaoli C. Microstructural and physiological features of tissues elucidated by quantitative- diffusion-tensor MRI. J Magn Reson B. 1996 Jun;111(3):209–19.

5. Radua J, Vieta E, Shinohara R, Kochunov P, Quidé Y, Green MJ, et al. Increased power by harmonizing structural MRI site differences with the ComBat batch adjustment method in ENIGMA. NeuroImage. 2020 Sep;218:116956.

6. Dansereau C, Benhajali Y, Risterucci C, Pich EM, Orban P, Arnold D, et al. Statistical power and prediction accuracy in multisite resting-state fMRI connectivity. NeuroImage. 2017 Apr 1;149:220–32.

7. Radabaugh HL, Harris NG, Wanner IB, Burns MP, McCabe JT, Korotcov AV, et al. Translational Outcomes Project in Neurotrauma (TOP-NT) Pre-Clinical Consortium Study: A Synopsis. J Neurotrauma. 2025 Jan 22;

8. Kamnaksh A, Puhakka N, Ali I, Smith G, Aniceto R, McCullough J, et al. Harmonization of pipeline for preclinical multicenter plasma protein and miRNA biomarker discovery in a rat model of post-traumatic epileptogenesis. Epilepsy Res. 2019 Jan 1;149:92–101.

9. Magnotta VA, Matsui JT, Liu D, Johnson HJ, Long JD, Bolster BD, et al. Multicenter reliability of diffusion tensor imaging. Brain Connect. 2012;2(6):345–55.

10. Wilkinson MD, Dumontier M, Aalbersberg IjJ, Appleton G, Axton M, Baak A, et al. The FAIR Guiding Principles for scientific data management and stewardship. Sci Data. 2016 Mar 15;3(1):160018.

11. Hu F, Chen AA, Horng H, Bashyam V, Davatzikos C, Alexander-Bloch A, et al. Image harmonization: A review of statistical and deep learning methods for removing batch effects and evaluation metrics for effective harmonization. NeuroImage. 2023 Jul 1;274:120125.

12. Fortin JP, Parker D, Tunç B, Watanabe T, Elliott MA, Ruparel K, et al. Harmonization of multi-site diffusion tensor imaging data. NeuroImage. 2017 Nov 1;161:149–70.

13. Johnson WE, Li C, Rabinovic A. Adjusting batch effects in microarray expression data using empirical Bayes methods. Biostatistics. 2007 Jan 1;8(1):118–27.

14. Wengler K, Cassidy C, van der Pluijm M, Weinstein JJ, Abi-Dargham A, van de Giessen E, et al. Cross- Scanner Harmonization of Neuromelanin-Sensitive MRI for Multisite Studies. J Magn Reson Imaging. 2021;54(4):1189–99.

15. Bell TK, Godfrey KJ, Ware AL, Yeates KO, Harris AD. Harmonization of multi-site MRS data with ComBat. NeuroImage. 2022 Aug 15;257:119330.

16. Richter S, Winzeck S, Correia MM, Kornaropoulos EN, Manktelow A, Outtrim J, et al. Validation of cross- sectional and longitudinal ComBat harmonization methods for magnetic resonance imaging data on a travelling subject cohort. Neuroimage Rep. 2022 Dec;2(4):None.

17. Quach M, Ali I, Shultz SR, Casillas-Espinosa PM, Hudson MR, Jones NC, et al. ComBating inter-site differences in field strength: harmonizing preclinical traumatic brain injury MRI data. NMR Biomed. 2024 Aug;37(8):e5142.

18. Wanner IB, McCabe JT, Huie JR, Harris NG, Paydar A, McMann-Chapman C, et al. Prospective Harmonization, Common Data Elements, and Sharing Strategies for Multicenter Pre-Clinical Traumatic Brain Injury Research in the Translational Outcomes Project in Neurotrauma Consortium. J Neurotrauma. 2025 Jan 20;

19. Sert NP du, Hurst V, Ahluwalia A, Alam S, Avey MT, Baker M, et al. The ARRIVE guidelines 2.0: Updated guidelines for reporting animal research. PLOS Biol. 2020 Jul 14;18(7):e3000410.

20. Lee SH, Ban W, Shih YYI. BrkRaw/bruker: BrkRaw v0.3.3 [Internet]. Zenodo; 2020 [cited 2025 Mar 31]. Available from: https://zenodo.org/records/3877179

21. Smith SM, Zhang Y, Jenkinson M, Chen J, Matthews PM, Federico A, et al. Accurate, robust, and automated longitudinal and cross-sectional brain change analysis. NeuroImage. 2002 Sep;17(1):479–89.

22. Salimi-Khorshidi G, Douaud G, Beckmann CF, Glasser MF, Griffanti L, Smith SM. Automatic denoising of functional MRI data: combining independent component analysis and hierarchical fusion of classifiers. NeuroImage. 2014 Apr 15;90:449–68.

23. Kellner E, Dhital B, Kiselev VG, Reisert M. Gibbs-ringing artifact removal based on local subvoxel-shifts. Magn Reson Med. 2016;76(5):1574–81.

24. Andersson JLR, Skare S, Ashburner J. How to correct susceptibility distortions in spin-echo echo-planar images: application to diffusion tensor imaging. NeuroImage. 2003 Oct;20(2):870–88.

25. Smith SM, Jenkinson M, Woolrich MW, Beckmann CF, Behrens TEJ, Johansen-Berg H, et al. Advances in functional and structural MR image analysis and implementation as FSL. NeuroImage. 2004;23 Suppl 1:S208–219.

26. Woolrich MW, Jbabdi S, Patenaude B, Chappell M, Makni S, Behrens T, et al. Bayesian analysis of neuroimaging data in FSL. NeuroImage. 2009 Mar 1;45(1, Supplement 1):S173–86.

27. Tournier JD, Smith R, Raffelt D, Tabbara R, Dhollander T, Pietsch M, et al. MRtrix3: A fast, flexible and open software framework for medical image processing and visualisation. NeuroImage. 2019 Nov 15;202:116137.

28. Avants BB, Tustison NJ, Song G, Cook PA, Klein A, Gee JC. A reproducible evaluation of ANTs similarity metric performance in brain image registration. NeuroImage. 2011 Feb 1;54(3):2033–44.

29. Mayer AR, Bedrick EJ, Ling JM, Toulouse T, Dodd A. Methods for identifying subject-specific abnormalities in neuroimaging data. Hum Brain Mapp. 2014 Nov;35(11):5457–70.

30. Verley DR, Torolira D, Pulido B, Gutman B, Bragin A, Mayer A, et al. Remote Changes in Cortical Excitability after Experimental Traumatic Brain Injury and Functional Reorganization. J Neurotrauma. 2018 Oct 15;35(20):2448–61.

31. Paydar A, Harris NG. The pericontused cortex can support function early after TBI but it remains functionally isolated from normal afferent input. Exp Neurol. 2023 Jan 1;359:114260.

32. Avants BB, Tustison NJ, Stauffer M, Song G, Wu B, Gee JC. The Insight ToolKit image registration framework. Front Neuroinformatics. 2014;8:44.

33. Whitcher B, Schmid VJ, Thorton A. Working with the DICOM and NIfTI Data Standards in R. J Stat Softw. 2011 Oct 27;44:1–29.

34. Muschelli J. neurobase: “Neuroconductor” Base Package with Helper Functions for “nifti” Objects [Internet]. 2024 [cited 2025 Mar 31]. Available from: https://cran.r-project.org/web/packages/neurobase/

35. Wickham H. ggplot2: Elegant Graphics for Data Analysis [Internet]. New York, NY: Springer; 2009 [cited 2025 Mar 31]. Available from: https://link.springer.com/10.1007/978-0-387-98141-3

36. McCarthy P. FSLeyes [Internet]. Zenodo; 2024 [cited 2025 Mar 30]. Available from: https://zenodo.org/records/11047709

37. Ho J, Tumkaya T, Aryal S, Choi H, Claridge-Chang A. Moving beyond P values: data analysis with estimation graphics. Nat Methods. 2019 Jul;16(7):565–6.

38. Champely S. pwr: Basic Functions for Power Analysis [Internet]. 2006 [cited 2025 Mar 30]. p. 1.3-0. Available from: https://CRAN.R-project.org/package=pwr

39. Han Q, Xiao X, Wang S, Qin W, Yu C, Liang M. Characterization of the effects of outliers on ComBat harmonization for removing inter-site data heterogeneity in multisite neuroimaging studies. Front Neurosci. 2023;17:1146175.

40. Yu M, Linn KA, Cook PA, Phillips ML, McInnis M, Fava M, et al. Statistical harmonization corrects site effects in functional connectivity measurements from multi-site fMRI data. Hum Brain Mapp. 2018 Nov;39(11):4213–27.

41. Horng H, Singh A, Yousefi B, Cohen EA, Haghighi B, Katz S, et al. Improved generalized ComBat methods for harmonization of radiomic features. Sci Rep. 2022 Nov 8;12(1):19009.

42. Smith G, Santana-Gomez CE, Staba R, Harris NG. Unbiased Population-Based Statistics to Obtain Pathologic Burden of Injury after Experimental TBI [Internet]. bioRxiv; 2025 [cited 2025 Apr 8]. p. 2025.04.03.647083. Available from: https://www.biorxiv.org/content/10.1101/2025.04.03.647083v1

43. Button KS, Ioannidis JPA, Mokrysz C, Nosek BA, Flint J, Robinson ESJ, et al. Power failure: why small sample size undermines the reliability of neuroscience. Nat Rev Neurosci. 2013 May;14(5):365–76.

44. Harris NG, Verley DR, Gutman BA, Sutton RL. Bi-directional changes in fractional anisotropy after experiment TBI: Disorganization and reorganization? Neuroimage. 2016 Jun;133:129–43.

